# FishNET: An automated relational database for zebrafish colony management

**DOI:** 10.1101/494971

**Authors:** Abiud Cantu Gutierrez, Manuel Cantu Gutierrez, Alexander M. Rhyner, Oscar E. Ruiz, George T. Eisenhoffer, Joshua D. Wythe

**Author notes:** To whom correspondence should be addressed: Joshua D. Wythe CVRI, Department of Molecular Physiology and Biophysics, Baylor College of Medicine, One Baylor Plaza, Houston, TX 77030.

## Abstract

The zebrafish *Danio rerio* is a powerful model system to study the genetics of development and disease. However, maintenance of zebrafish husbandry records is both time intensive and laborious, and a standardized way to manage and track the large amount of unique lines in a given laboratory or centralized facility has not been embraced by the field. Here we present FishNet, an intuitive, open source, relational database for managing data and information related to zebrafish husbandry and maintenance. By creating a “virtual facility”, FishNET enables users to remotely inspect the rooms, racks, tanks and lines within a given facility. Importantly, FishNET scales from one laboratory, to an entire facility with several laboratories, to multiple facilities, generating a cohesive laboratory and community-based platform. Automated data entry eliminates confusion regarding line nomenclature and streamlines maintenance of individual lines, while flexible query forms allow researchers to retrieve database records based on user-defined criteria. FishNet also links associated embryonic and adult biological samples with data, such as genotyping results or confocal images, to enable robust and efficient colony management and storage of laboratory information. A shared calendar function with email notifications and automated reminders for line turnover, automated tank counts and census reports promote communication with both end-users and administrators. The expected benefits of FishNET are improved vivaria efficiency, increased quality control for experimental numbers, and flexible data reporting and retrieval. FishNet’s easy, intuitive record management and open source, end user-modifiable architecture provides an efficient solution to real-time zebrafish colony management for users throughout a facility and institution, and in some cases across entire research hubs.

## INTRODUCTION

The fecundity, rapid development, external fertilization, amenability to both forward^1-7^ and reverse^8-11^ genetic approaches, conservation of core vertebrate protein-coding genes^12^, small size and inexpensive husbandry costs make zebrafish a powerful vertebrate model for studying human physiology and disease^13,14^. Moreover, their optical transparency^15-17^ and readily available suite of genetic tools (Tol tranposases, Gal4/UAS, Cre/loxP, etc.)^18-27^ and fluorescent reporters (e.g. gCaMP, lifeact)^18,28^ have made zebrafish the premier model system for studying vertebrate biology in real-time. These features, combined with their impressive regenerative capacity, also make zebrafish ideal for studying vertebrate tissue and organ regeneration^29,30^. Given these advantages, it is no wonder that fields as diverse as developmental neurobiology^31^ and cancer^32^ have leveraged the embryonic and adult zebrafish, respectively, to make valuable biological insights. As more labs have embraced this model system, there has been an explosion in the amount of transgenic and mutant lines generated by the community, and shared use zebrafish “core” facilities continue to be built across the country, all creating a significant need for accurate, automated, real-time colony management tracking software within laboratory groups, throughout a facility and institution, and in some cases across entire research hubs.

Despite the importance of effective data management in animal research, many investigators employ handwritten notebooks or spreadsheet applications for managing small and medium sized animal colonies. While these *ad hoc* data entry approaches offer the benefit of being simple to adopt, they do not scale with increased user numbers (even within a single laboratory) and they are not practical across an entire facility. Relying upon individuals to enter the correct line nomenclature (e.g. allele) and accurate information (e.g. sex, date of birth, etc.), unnecessarily exposes these “databases” to avoidable operator error. Additionally, paper records have the significant drawback that they cannot be accessed simultaneously by multiple users, they cannot be viewed remotely, and they can be misplaced or destroyed. On the other hand, electronic spreadsheet databases suffer from their own limitations, as multiple users cannot access them simultaneously in real-time. In addition, data stored in this format are challenging to mine, and they typically lack a uniform/controlled lexicon and best practice rules for data entry are absent, thus making them highly susceptible to operator error. Moreover, simple spreadsheets (e.g. Excel) cannot navigate complex data structures, such as breeding schemes or pedigrees, as required in a model colony management software.

Paraphrasing Silver^33^, and as epitomized in the Jax Colony Management Software for mice^34^, an ideal database would track (1) individual animals and their ancestors; (2) matings between animals; (3) progeny born from such matings (e.g litters) and the individuals within litters that are used in experiments; and (4) experimental materials (tissues and DNA samples) obtained from individual animals. Such a database should also format records so that determination of relationships between any components of the colony is possible (e.g. which tissue came from which animal). Ideally, this platform would feature an intuitive graphical user interface ^12^, have a robust online community to support trouble shooting or modifications to the underlying architecture or functionality, and also be open source.

A handful of software solutions specifically designed for managing zebrafish animal husbandry exist. A summary of some of the currently available zebrafish colony management applications, both commercial and open source, is provided in **Table 1**, while their functionalities are summarized in **Table 2.** Available solutions for data management also vary significantly in terms of cost, implementation requirements, flexibility, and level of user support. Additionally, few of these options are open source, and even fewer also record phenotyping and experimental data. These drawbacks, along with the cost of commercial options, or the potentially intimidating learning curve associated with adopting open source options – such as creation and maintenance of SQL databases, along with knowledge of programming languages like Python or PHP to manage them – may explain the reluctance of the zebrafish community to embrace these solutions.

**Table 1.**
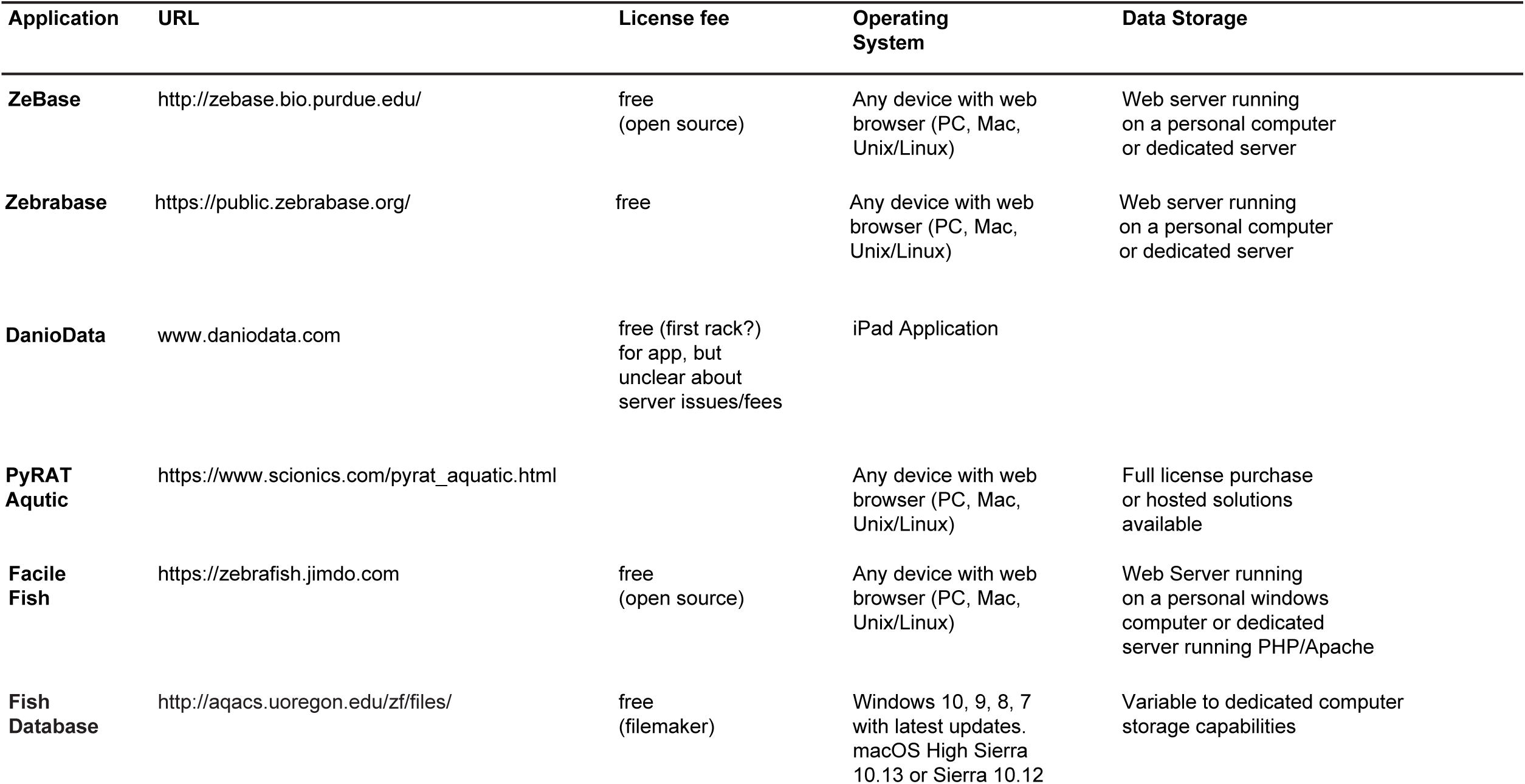
Examples of currently available zebrafish colony management applications.

| Application | URL | License fee | Operating System | Data Storage |
| --- | --- | --- | --- | --- |
| **ZeBase** | http://zebase.bio.purdue.edu/ | free<br>(open source) | Any device with web browser (PC, Mac, Unix/Linux) | Web server running on a personal computer or dedicated server |
| **Zebrabase** | https://public.zebrabase.org/ | free | Any device with web browser (PC, Mac, Unix/Linux) | Web server running on a personal computer or dedicated server |
| **DanioData** | www.daniodata.com | free (first rack?) for app, but unclear about server issues/fees | iPad Application |  |
| **PyRAT Aquatic** | https://www.scionics.com/pyrat_aquatic.html |  | Any device with web browser (PC, Mac, Unix/Linux) | Full license purchase or hosted solutions available |
| **Facile Fish** | https://zebrafish.jimdo.com | free<br>(open source) | Any device with web browser (PC, Mac, Unix/Linux) | Web Server running on a personal windows computer or dedicated server running PHP/Apache |
| **Fish Database** | http://aqacs.uoregon.edu/zf/files/ | free<br>(filemaker) | Windows 10, 9, 8, 7 with latest updates.<br>macOS High Sierra 10.13 or Sierra 10.12 | Variable to dedicated computer storage capabilities |

**Table 2.**
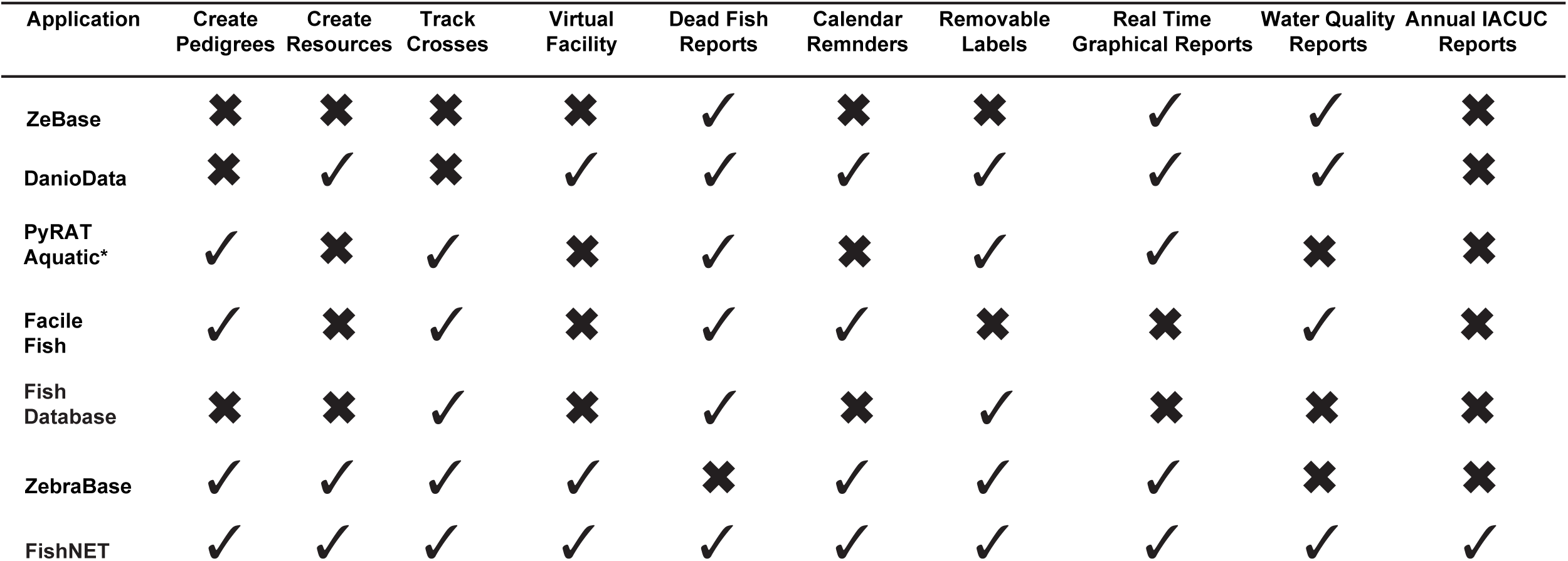
Features of currently available zebrafish colony management applications.

Here, we describe FishNET. FishNET is a comprehensive relational database application developed specifically to meet biomedical research community demands for a well-engineered, flexible database system supporting zebrafish animal husbandry and data management. FishNET meets the aforementioned criteria for an ideal record keeping system, with several added benefits: including, a remotely accessible virtual facility view; a pedigree function; barcode scannable labels for tanks, crosses, embryos, and fry; a calendar function that emails users for events such as line turnover or graduation of fry to the nursery; sick fish reports; records for genotyping protocols; and real-time records of all fish tanks, their use, and water quality in a facility for billing purposes and IACUC reporting purposes.

FishNET runs directly on Macintosh or PC computers using the FileMaker Pro Advanced (FMPA) application. Hosted databases can also be viewed remotely via web browsers on PC, Macintosh, or Linux computers by incorporating Filemaker WebDirect. Furthermore, it can be accessed on iPad and iPhones through the Filemaker Go application in the App Store (https://itunes.apple.com/us/genre/ios/id36?mt=8). The functional features of FishNET include support for controlled vocabularies (such as uniform allele nomenclature to avoid confusion regarding the provenance of a transgenic or mutant line), multiuser capabilities, quick and accurate data reporting, pedigree tracking, animal husbandry workflow, sample tracking, and experimental data capture. The expected benefits of FishNET are improved vivaria efficiency, increased quality control for experimental numbers, and flexible data reporting and retrieval. FishNET is a freely available tool, and FileMaker is a well-supported application with a large user base. Free trial versions of FileMaker Pro Advanced 17 are available at https://content.filemaker.com/filemaker-trial-en-rf. FileMaker Go (for iOS mobile devices) is available at the App Store (https://www.filemaker.com/products/filemaker-go/). A full database, as well as an empty shell, of FishNET are freely available at http://www.wythelab.com/wythe-lab-databases.

## MATERIALS AND METHODS

### System Architecture

FishNET employs a normalized relational database that can be run through FMPA. The relational data model that underlies FishNET minimizes data redundancy, enforces data integrity, leverages controlled vocabularies to ensure proper allele designations, and enables accurate data retrieval against large data sets. The user interface enables addition of new functionality and enhancement of existing functions without major modifications to the basic infrastructure, enabling easy adaptation to user-specific needs. FMPA ensures safe multi-user concurrent data entry and editing in real-time. Additionally, FishNET features a built-in barcode generator and barcode reader, and when these features are combined with a validated printer (as we demonstrate herein) together provide instant recognition and tracking of all fish stock information and related actions on a mobile device.

An overview of the two primary configurations of FishNET is shown in **Figure 1A**. FishNET can be set up to run as a stand-alone database on a personnel computer running a Windows or MacOS operating system. Alternatively, FishNET can be hosted on a central computer running a Macintosh or Windows operating system using FMPA Server software to support small-to medium-sized research groups where FMPA client software is installed on all user desktop and laptop computers (e.g. 1 license of FMPA per accessing computer). This same FMPA Server configuration also supports use of FishNET through the FileMaker Go App on mobile handheld devices with barcode readers running iOS 11.2 or later (e.g. iPad, iPhone). Handheld computers that have been tested include iPad Air 2 as well as iPhone 6s and iPhone X. In the server configuration, FishNET can be accessed – with appropriate user authentication capabilities – on any internet capable device running Safari, Chrome, Internet Explorer, or Microsoft Edge internet browsers using FileMaker WebDirect.

**Figure 1.**
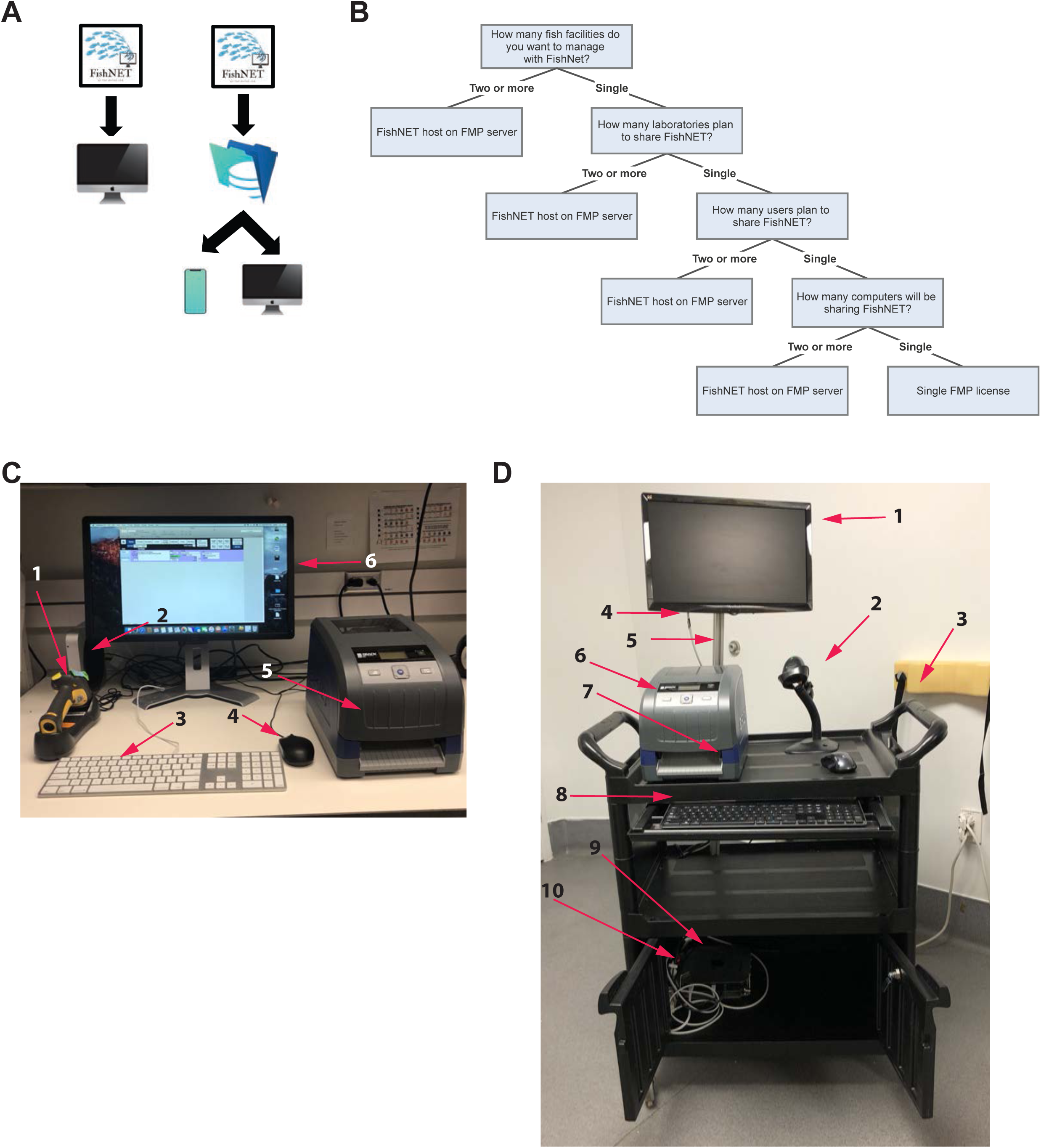
Schematic representation of FishNET Configuration. A) FishNET can be used as a single unit installed in one PC alone or through a server that can support multiple computers and handheld devices. B) Decision tree to decide if a single FMPA License is needed, or multiple licenses hosted on a server are required. C) The stationary, standalone desktop configuration for running FishNET consists of: 1. Barcode Scanner. 2. Mac Mini. 3. Keyboard. 4. Mouse. 5. BBP33 Brady printer. 6. Computer monitor. D) The mobile configuration for FishNET consists of: 1. 19” Widescreen LED Backlight LCD Monitor. 2. Barcode Scanner. 3. Dual-Handle Plastic Enclosed Shelf Service Cart. 4. DisplayPort to VGA Adapter Converter Cable. 5. V-Slot Extruded Aluminum. 6. BBP33 Brady printer. 7. Mac Mini with Mac Mini Security Mount.

The cost of setting up and hosting FishNET will vary depending on the configuration. The number of users and their need to modify the underlying database architecture (e.g. do they need to function as an administrator, or simply upload, access, and download data) will dictate whether a standalone license is sufficient, or multiple licenses that each connect to a FMPA Server (hosted via an on-premise server) or FileMaker Cloud (hosted remotely via FileMaker, not described here) are required. At this time, an annual FMPA license costs $540 for 5 licenses per year (academic pricing) with FMPA Server included, while a permanent individual instance of FMPA costs $324, and FileMaker Cloud costs $400 (user licenses not included). The cost to host FMP Server will vary by institution and will also depend on whether the institution hosts it on their own server (which requires Windows Server 2016, or newer) or if it is instead hosted locally within the laboratory (for instance, on a computer running MacOS Sierra 10.12 (or newer) software). A guide to determining which license is appropriate for your group is provided in **Figure 1B**, as are images of the station set up in **Figure 1C** and **1D**. A more detailed comparison between the costs of FMP Server and Cloud is provided in **Supplemental Table 1**.

### Hardware Requirements

The base hardware requirements of FileMaker Pro Advanced and FishNET are relatively modest. FishNET runs smoothly on a late 2014 Mac mini with a 2.6 GHz, dual-core Intel i5 (2278U) processor, with 8GB of 1600 MHz LPDDR3 onboard memory, with 3MB virtual memory, and a 1 TB hard drive and 802.11ac Wi-Fi wireless networking and Bluetooth 4.0 wireless technology requiring 100-240V AC. While the database itself can be set up multiple ways, below we describe two standard configurations. One option is an immobile station on a bench top within the facility room that requires a wireless barcode reader integrated with the computer (to read bar codes, enter data) as well as a label printer (for the racks, tanks, crosses, and fry) (**Figure 1C**). For multiple users, a more sensible configuration may be to locally host FishNET (this requires a FileMaker Pro Advanced Server license, which is included in the purchase of a team FMPA license – which covers 5 or more individual licenses). This set up allows for simultaneous, multi-user access and data entry using either FMPA on the same network, or through an internet web browser on the same network via FileMaker WebDirect. In this configuration access on the same network could also be achieved on a handheld device using the FileMaker Go application (which is restricted to iPads and iPhones running iOS 11.2 or above). Individual licenses for FMPA would only be required for users that want to easily print labels with barcodes, run the automated genotyping annotation script (see below), or those that need to modify the underlying database architecture. The costs for this configuration are outlined in **Table 3**.

**Table 3.** Hardware required for the immobile, desktop FishNET configuration.

| Item | Use | Cat # | Manufacturer | Cost (Vendor) |
| --- | --- | --- | --- | --- |
| **Mac Mini Security Mount** | To immobilize the Mac Mini computer/security | MMEN76 | Maclocks.com | \$49.95<br>(Maclocks.com) |
| **Mac Mini** | Computer | MGEN2LL/A | Apple | \$699 (Apple) |
| **19" Widescreen LED Backlight LCD Monitor** | Visualization | VA1938WA-LED | ViewSonic | \$102 (CDW) |
| **6 outlet, 6 ft, Power strip with surge protector** | Power | Super 7 Protect it! Surge protector S9TLP606 | Tripp Lite | \$14.40<br>(officemax) |
| **Bluetooth, Waterproof Keyboard** | Data entry | WKB-2000BW (for Mac) | Adesso | \$29.95<br>(Amazon) |
| **Wireless Mouse** | Data entry, navigation | M510 mouse, 910-001822 | Logitech | \$38.99<br>(CDW) |
| **Wireless, Waterproof Barcode scanner** | Barcode scanning, data entry | 19 140 592 | Adesso | \$125<br>(newegg.com) |
| **BBP33 W/AUTO CUT PRINTER** | Print Labels | 9SIAFJ86WS6511 | Brady | \$1,169.33<br>(Fisher Scientific) |
| **RIBBON, R6200 BLK, 4.33 X200FT** | Ribbon needed to print labels | 19 140 590 | Brady | \$48.72<br>(Fisher Scientific) |
| **Direct Custom Labels 1,000** | Cling labels | NC1568023 | Brady | \$469.26<br>(Fisher Scientific) |
| **Optional: iPad 32 GB, Wi-Fi Only, 5th Generation** | Data entry, visualization | MP2F2LL/A | Apple | \$279<br>(Target) |

A second configuration is for a mobile station that can travel throughout an entire facility (**Figure 1D**). The costs and components of this set up are shown in **Table 4**.

**Table 4.** Hardware required for the mobile FishNET configuration.

| Application | Use | Cat # | Manufacturer | Cost (Vendor) |
| --- | --- | --- | --- | --- |
| Dual-Handle Plastic Enclosed Shelf Service Cart | For Mac Mini and peripherals | #940Z122 | Gamut | \$447.19 (Gamut.com) |
| Backup Portable Battery | Battery for Mobility | APC-Back UPS PRO Pro BR1000G | APC | \$133 (Amazon) |
| Rocketfish 13-26" Tilting Low-profile TV Wall Mount | To mount PC monitor to aluminum rod | Rf-tvmlpto1v2 | Rocketfish | \$21 (ebay) |
| V-SLOT Extruded Aluminum 20X20X1500 | To hold PC monitor to cart | AAA-019 | Hobby-Fab | \$17.69 (ebay) |
| 6 ft Mini DisplayPort to VGA Adapter Converter Cable – mDP to VGA 1920x1200 | To wire Mac Mini to PC monitor | MDP2VGAMM6W | StarTech | \$41.99 (StarTech.com) |

### Software Requirements

Regardless of the hardware set up, the main drawback of using a commercial relational database platform is that while it is open source, non-coding based, and user modifiable, a facility or laboratory must purchase a permanent or annual license to use FileMaker Pro Advanced. Fortunately, several flexible options exist, as outlined below. Additionally, the FileMaker Go App is compatible with Apple mobile devices (e.g. iPhone and iPad). If FishNET will be hosted, then a computer with a Dual Core CPU processor, 8GB of RAM, and at least 80 GB of hard drive storage, running either Windows Server 2016 Standard Edition (with Desktop Experience), Windows Server 2012 R2 with Update, macOS High Sierra 10.13 or macOS Sierra 10.12 will be required. The FMP Server license is included when you obtain at least 5 licenses of FMPA, making this the logical choice for most user groups.

### Data Security and Backup

To protect the user’s data, FMPA Server offers AES-256 encryption. It also supports third-party, secure sockets layer ^17^ certificates to establish secure links between the server and web browsers. Furthermore, using an institution’s wireless local area network (Wi-Fi) to transfer files from the FMPA Server to individual users adds another layer of security (provided that the institution requires user authentication to connect to the network/has a secure firewall).

A centralized database, such as FishNET, offers the advantage of simplifying data redundancy and mirroring (e.g. creating backups). When using a single instance of FMPA on a single computer, a user or lab can back up the entire contents of the hard drive, along with the FMPA database file(s), using a standard backup program (e.g. Time Machine, Super Duper, etc.), storing these copies on an external hard drive or remotely in the cloud. In the case of FMPA Server, the program by default creates a backup daily, storing the last seven backups (and successively re-writing over them from oldest to newest), with the option to keep specific backups at will. In this case, we suggest copying FishNET to an external drive.

## RESULTS

### General Overview, Installing the Database

We have developed a system for positional identification of tanks and lines based on user-defined facilities, rooms, racks, tanks, and lines. The underlying infrastructure of this system is a relational database. A typical database is composed of tables of individual entries, or *records* (as they are referred to in FMPA), that contain data (in the case of FishNET these data are information such tank number, genotype, date of birth). In a relational database these records relate to one another through shared or common data (e.g. tank identifier number, genotype, etc.) and as such these records (and all of their associated data) can be sorted, queried, and viewed independently, or in a list format. Tables of records within FishNET can be broadly grouped into seven categories (or *layouts*, in FMPA terms): Tanks, Crosses, Lines, Harvests, Nursery, Labs, and Statistics. Information regarding the allele (Lines), parental strain(s)/ mating history (Tanks and Crosses), progeny (nursery), owner of the tank/IACUC number(s) associated with a tank (Labs and Tanks), usage (Harvests), and water quality/births/deaths (Statistics) are all stored in these various records, enabling robust and detailed records of zebrafish husbandry within a research group or across an entire facility. An eighth category, Virtual Facility, which contains unique identifies for the facility, room, rack, row, and column of a given tank, creates a virtual facility containing room(s), rack(s), tank(s), and nursery of unique zebrafish lines, thus enabling users or administrators to view all occupied tanks within the facility (as well as the pertinent information for each individual tank: sex, age, genotype) and easily and efficiently track and locate tanks within a facility. A final category is the Calendar layout, which relates to all other tables (e.g. crosses, tanks, nursery, etc.). This category has email functionality to create reminders for graduating fry to the main system or for turning over lines, or any other user-defined event. Finally, each tank, cross, and fry/larvae, as well as each rack, are uniquely barcoded to enable, fast, reliable data entry and functionality (such as moving tanks within or across racks, setting up matings, recording dead fish, etc.).

Once you have determined the correct FMPA configuration for your needs (e.g. locally hosted server, cloud, etc. (**Figure 1A-B**)), purchased and installed the software, visit www.wythelab.com/databases and download the latest version of FishNET. If using a single stand-alone FMPA license, the user only needs to open the FishNETv1.fmp12 file and proceed with the set up. If using the FMPA Server, the downloaded file needs to be placed in the FMP Server database folder. In macOS this folder is placed by default in /Library/FileMaker Server/Data/Databases or in Windows in C:/Program Files/FileMaker/FileMaker Server/Data/Databases. Below, we provide a logical, step-by-step guide for creating your virtual facility, entering in lines, and populating racks with these lines, an overview of which is provided in **Figure 2A**. An overview of all tables containing source data how they relate to each other can be found in **Figure 2B**.

**Figure 2.**
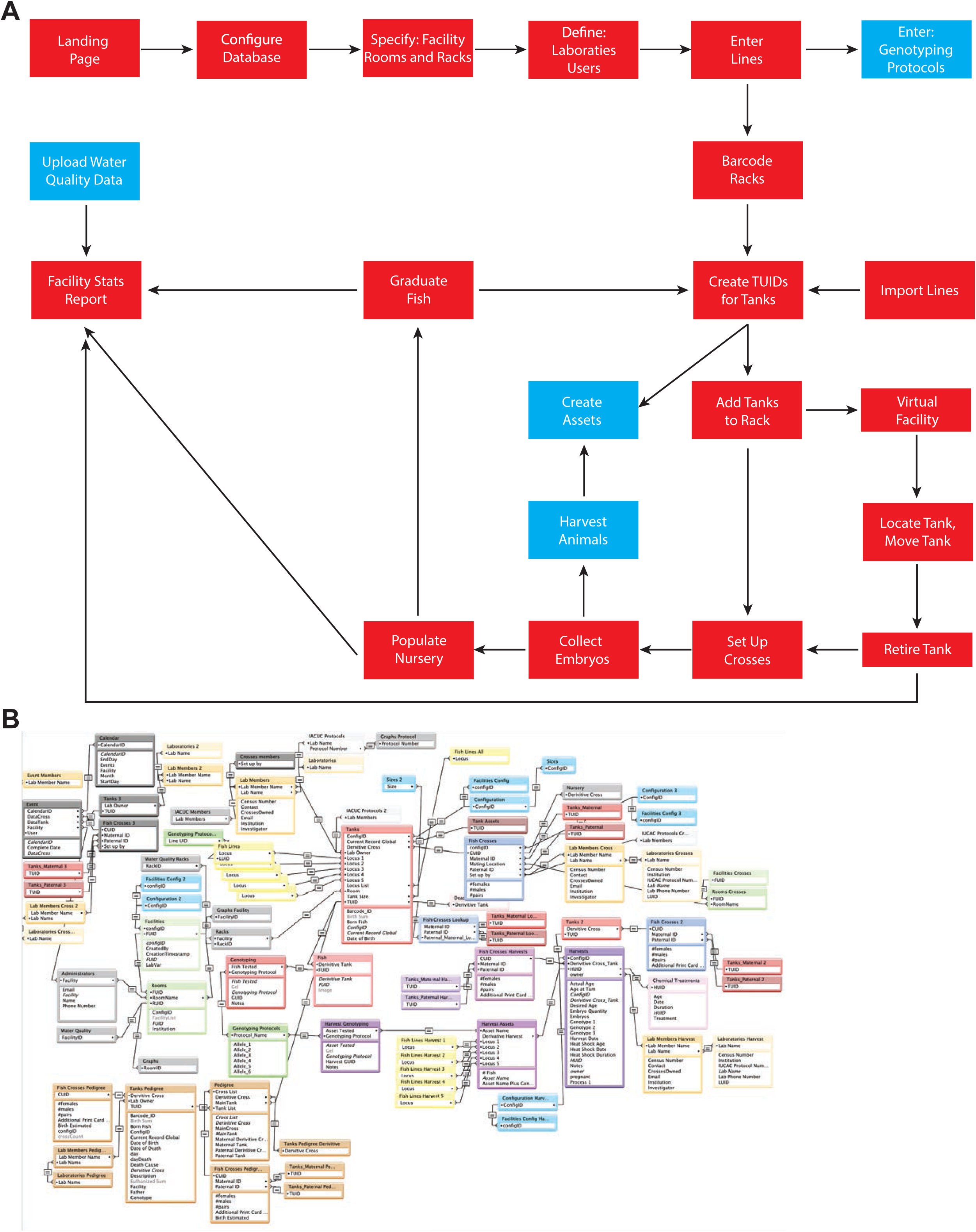
Step by step overview for implementing FishNET. A) The logic flow for configuring the database is shown, with essential steps in red and optional steps in blue. Simplified relationships are indicated by the directional arrows. B) Underlying relational view of FishNET showing all source tables that constitute the complete database.

### Importing FishNET to FMP

After depositing the downloaded file in the appropriate folder (as outline above), there needs to be one-time activation in the FileMaker Server Admin Console to bring FishNET online. This can be done by selecting the database and selecting “open” (**Figure 3A**). In order to add the database to individual FMPA instances, users need to select “Add App From Hosts”, then add the host using the IP Address of the server. This can be found in the FileMaker Server Admin Console (**Figure 3B**).

**Figure 3.**
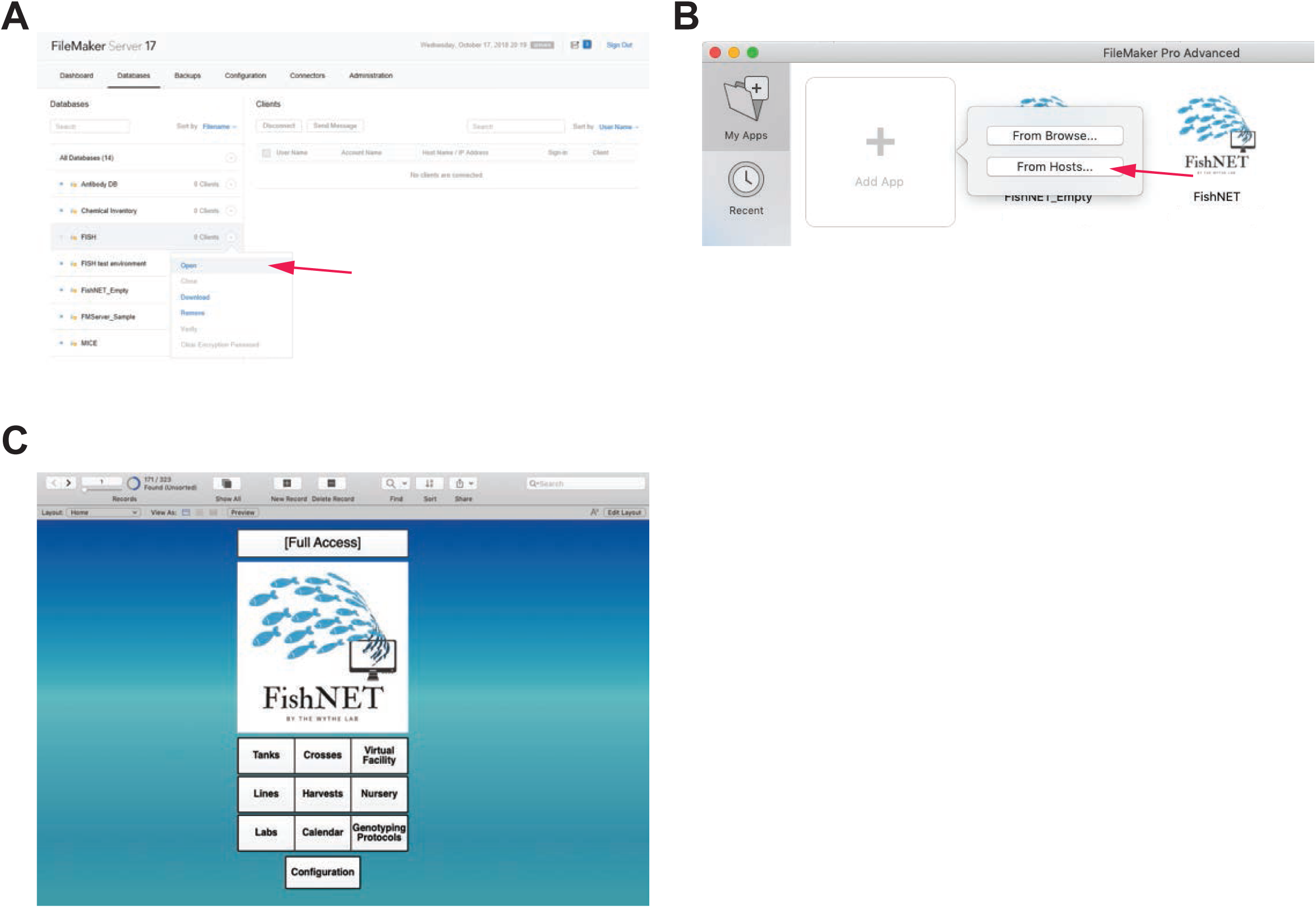
Importing FishNET to FileMaker Pro Advanced. A) Activation of FishNET database from the FileMaker Server 17 administrator console can be done by selecting “Open” (indicated by red arrow). B) FishNET App can be imported to FMP Advanced locally by selecting “From Browser…” or from FileMaker Server selecting “From Hosts…” (indicated by red arrow). C) FishNET landing page with full access privileges.

### Creating an Administrator User account

Upon opening the software, you should see the landing page (**Figure 3C**). Here, you will see a directory below the main FishNET icon. You can also select from the pulldown “layout” menu in the upper left-hand corner for navigation. From the “landing page”, or “home”, you will first select File / Manage… / Security. By default, an “admin user” is set up. If the database will be shared between different laboratories, we recommend setting up a password to access the administrator account, as this account can access all records in the database and can also change the underlying architecture of each layout, whereas the individual labs will only see and modify records that belong to their respective laboratories and are prevented from modifying database-wide records (**See Supplemental Video 1**). Creating labs and individual users will be addressed later.

### Creating Facilities, Rooms, and Racks

After generating an administrator profile, the next step is to configure facility (or facilities). This functionality is restricted to the administrator of the database, and not individual users. From the landing page, select the “Configuration” button. In this layout, in the first column the name of the facility needs to be entered (**Figure 4A**). In the second column, the different types of tanks being used are specified by size and the number of spaces they occupy on the rack(s). Following the same nomenclature, indicate the different tank capacities (for instance, if you are using a 5 Liter tank, enter **5L**) ordered from smallest (nursing tanks) to largest. The “Spaces Occupied” column is used as an approximation of the size of the tank in the zebrafish rack. **Figure 4B** shows an example set-up of a Tecniplast rack and the corresponding spaces occupied for tanks in each row. This information is used to create a customized virtual rack view. After this initial configuration has been created, the number of rooms within the facilities must be specified. This can be done by selecting “Virtual Facilities” from the landing page, upon clicking the button, a list of facilities should be shown in a pop up scroll down menu (**Figure 4C**). Selecting a facility will direct the user to the “facility view” layout. Here, the option to “add rooms” can be found (**Figure 4D**). Selecting “Virtual Facility” from the main menu bar can also direct the user to the rest of the facilities where rooms need to be set up. It is important to note that in order to use the virtual facility function, all facilities and rooms need to be properly labeled prior to inputting any records. This cannot be changed later without having to change all records linked to a room or facility. Finally, once the facility and number of rooms have been specified (and named), the number of racks per room must be entered by selecting the “view room” button (which takes users to view an individual room) (**Figure 4E**). When viewing an individual room, a user can select “Add Rack” to create digital racks that will reflect the physical room.

**Figure 4.**
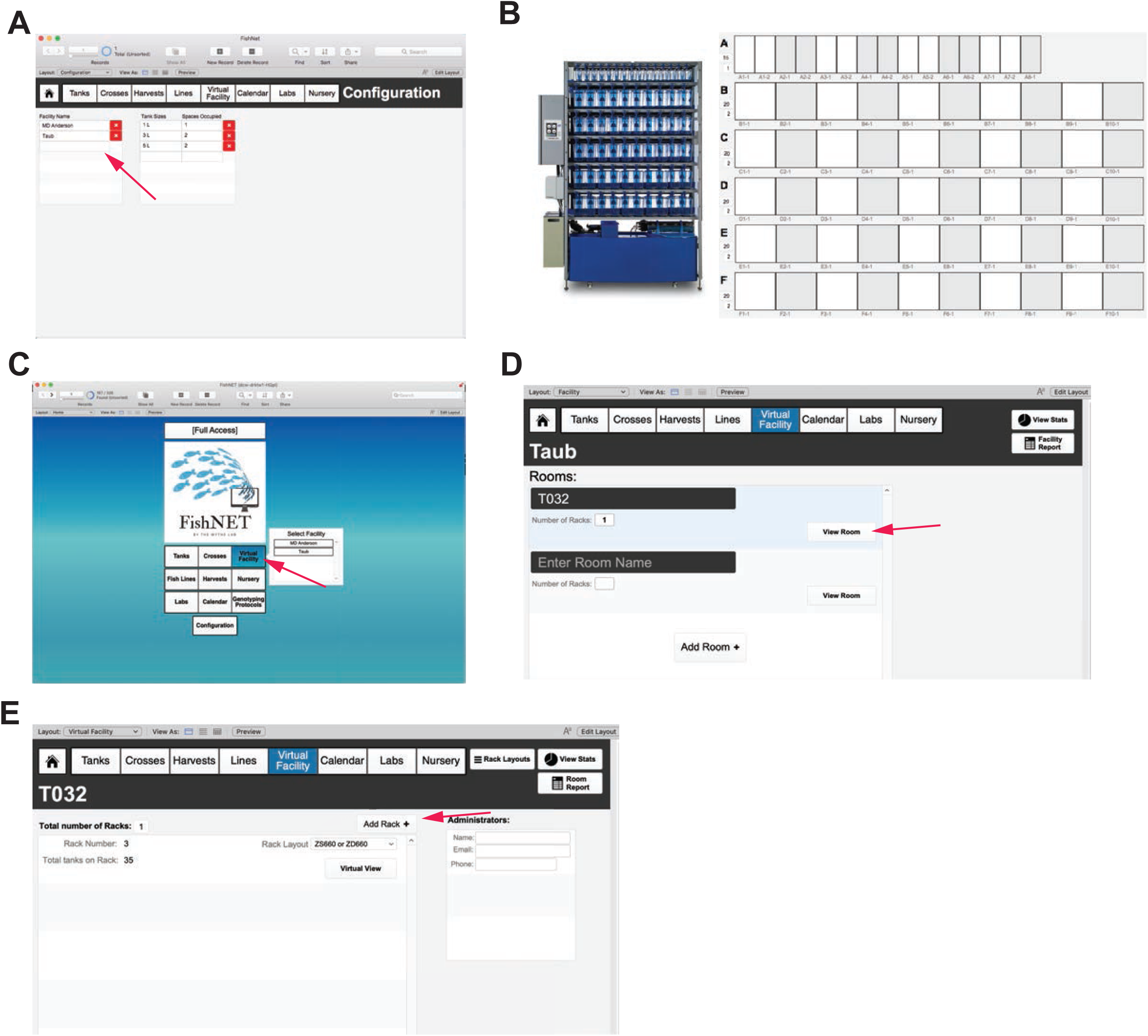
Initial configuration of facility, rooms and racks. A) Configuration page where facility names (indicated with red arrow), tank names, and spaces occupied are initially defined. This page should be set up before using FishNET and should not be modified later. B) A Tecniplast fish rack (on the left) with a corresponding virtual rack layout matching this configuration. C) After configuration, the virtual facilities button will now feature a dropdown list of facilities to choose from on the landing page (indicated with arrow). D) Within a facility, multiple separate rooms for housing zebrafish racks can be added by selecting “Add Room”. E) Inside each room, multiple fish racks can be added to house zebrafish, to add a rack simply select “Add Rack” (indicated with arrow).

For each rack that is added, the administrator will choose from predefined rack configurations corresponding to the current major commercial aquatic habitat manufacturers (e.g. Techniplast, Aquaneering, and Pentair) (**Figure 5A**). Pictures of the different aquatic systems can be viewed in the “Rack Layouts” section. Selecting one of these options creates a default layout of tank spaces matching these normal vendor configurations (**Figure 5B**). Upon selecting a rack layout, a virtual view of the rack will be shown, in the rack view, individual positions can be labeled with barcodes selecting the “Print Rack Labels” function (**Figure 5C**, indicated with red arrow). If the facility has a unique or custom rack layout, then select the “custom” option. To set up a custom rack configuration, select the number of tanks in each row (A, B, and so on) by entering in the top number, then specify the number of spaces they occupy by entering in the lower number (**Figure 5D**). Repeat this for each row on the rack. The “facility”, “room”, and “rack” categories provide spatial organization to the database to create a “virtual facility” that can be explored by the administrator, animal husbandry staff, and scientific users alike (more functions within this framework will be discussed after lines and tanks have been introduced). The categories of lines and tanks will be superimposed upon this user-defined framework later (**Supplemental Video 2**).

**Figure 5.**
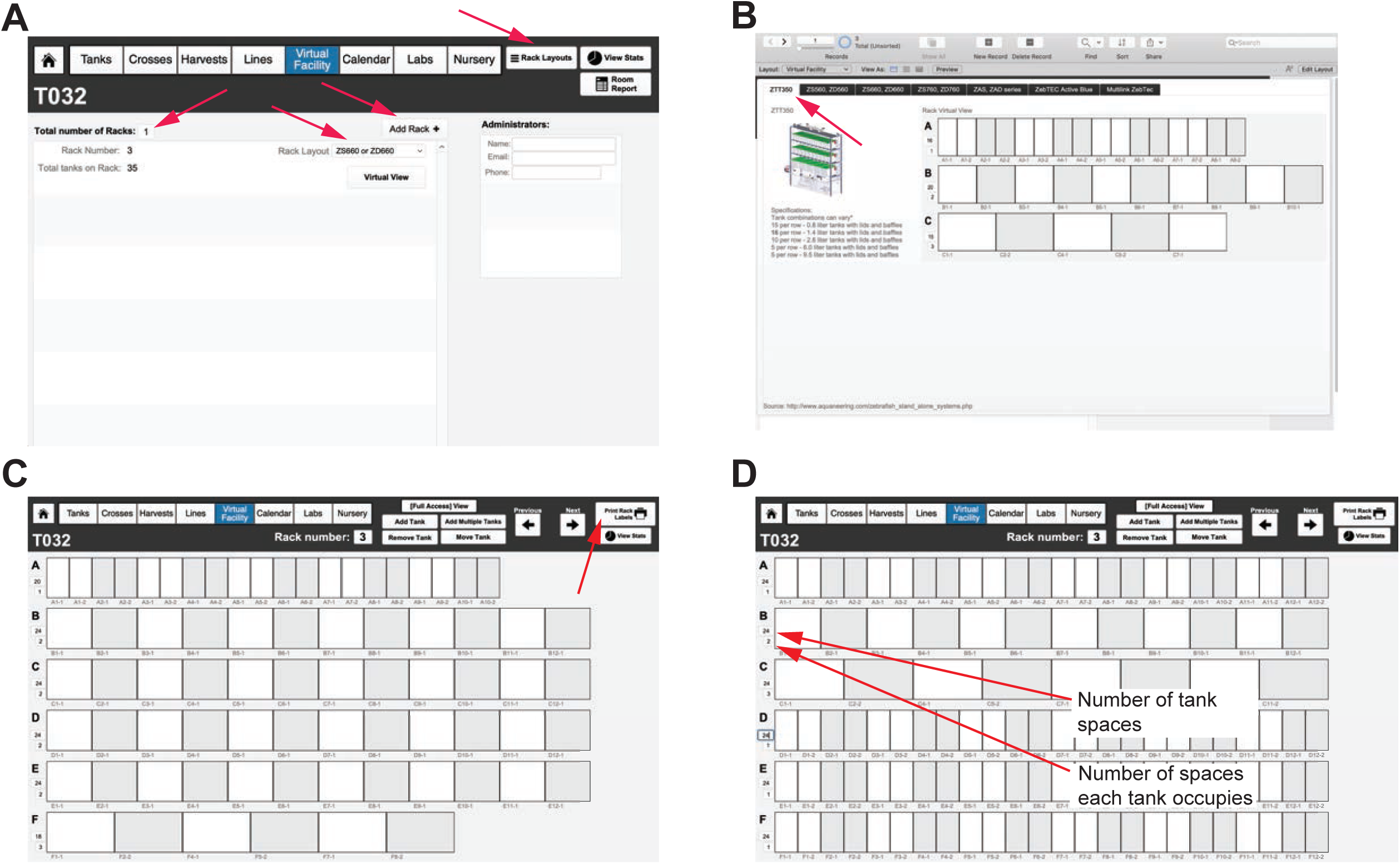
Creation of predefined and custom fish racks. A) Configuring the number and type of racks within a room of a facility. A drop-down list of predefined fish racks can be found, as well as pictures of each commercially available rack, by selecting the “Rack Layouts” button in the top, right corner of the menu header. Administrators for each room can also be added on the far right, main window. They can receive calendar notifications (fry to transfer to system, turnover of tanks, dead fish, etc.) via email, in addition to users. Racks are added using the “Add Rack” button and chosen from preconfigured options (that match commercially available racks) via the dropdown menu. B) A picture of a rack, and the corresponding virtual rack set up, for one of the predefined rack layouts (using the tabs at the top of the window). C) Virtual view of the newly set up, empty rack. Each space can be labeled with its own barcode using the “Print Rack Labels” function (indicated with red arrow). D) Example of a custom rack configuration. In each row, the number of tank spaces and how many spaces each tank occupies can be defined by the user. This can be changed to fit the custom rack specific layout by the facility administrator.

### Entering Laboratories and Individual Users

Next, individual laboratories and users within these laboratories will be entered into the database. Before beginning this section, have the IACUC/animal protocol number(s) for the lab, as well as the email address, phone number, and complete name of each user within a laboratory group readily available. The contact information for labs and users will later be assigned to individual tanks on the racks, as well as all crosses, and any fry to be raised, allowing for automated reminders about graduating fry to the nursery (or the main system), dead fish notifications, or providing details for contacting other users regarding available lines within the facility. To create a laboratory group, first select the “labs” button from the landing page (or dropdown layout menu). Once you are in the “laboratories” layout click the “+ New Record” button at the top of the FileMaker task bar (**Figure 6A**). Add all of the pertinent information to this empty field to create a laboratory. Once the lab has been entered, you can then populate it with users by selecting the “Lab Members” button. Within the “Lab Members” layout, click the “New Record” button to add users (**Figure 6B**). This information will later be used to establish ownership of individual tanks, crosses, and embryos within a facility. Finally, each lab should enter in a valid Institutional Animal Care and Use Committee (IACUC) protocol, and protocol duration. This information will be associated with all tanks, crosses, and actions of users within a laboratory (and present on all tank, cross, and nursery labels) (**Supplemental Video 3**).

**Figure 6.**
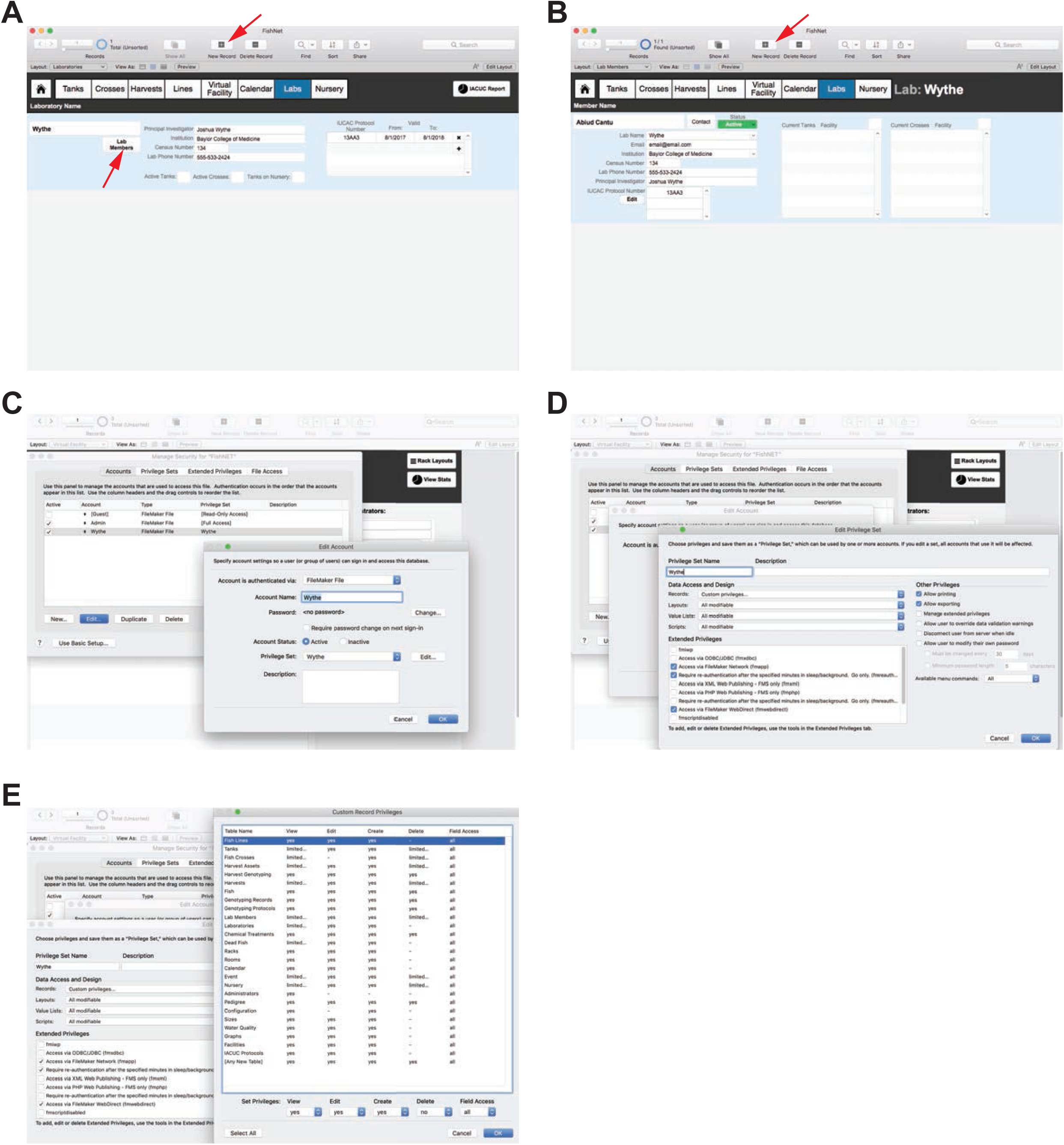
Entering laboratories and individual users. A) Example of a new laboratory, where the name of the lab, the PI, institution, contact info, and IACUC Protocol are all entered. Once lines and tanks have been entered, the lower part of this window will display the number of active tanks, active crosses, and tanks on the nursery for easy reference and billing purposes. Selecting the “Lab Members” button (indicated by the red arrow) will bring you to the next window. New laboratories can be added by selecting “+ New Record” option at the top of the FileMaker menu (indicated by a red arrow). B) Individual lab user set up. To add a new lab user, select the “+ New Record” option at the top of the FileMaker menu (indicated by a red arrow). Lab information is automatically populated from the general lab information window, and here individual contact details are entered. Users are defined as “active” or “inactive”. Later, tanks and crosses owned by this individual user will be shown in this window. C-E) After adding laboratory accounts, an administrator can set up privileges for all users in that lab, as well as other labs, to restrict or permit visualization of lines and handling of fish (e.g. setting up crosses, etc.) by non-lab members.

By default, all new laboratories have access to view and modify the complete database. Below are a series of permissions that an administrator can restrict if they wish to limit individual laboratory members’ access to certain features or functions within FishNET (otherwise, ignore the following steps). Specifically, the following commands will allow individual users to “see and modify” all information (e.g. genotype, number, sex, location) for tanks owned by all users in a common laboratory, but tanks from other laboratories within the facility (including administrators’ stock tanks), are hidden and the records locked for modification. Administrators, such as animal facility technicians, can view all information and record mortalities and health problems, and can work with any tank (e.g. to graduate fry or to retire a sick tank). For this type of access, first create a new user or group account (laboratory) with restricted access to the database, by selecting File / Manage / Security. Then select “New Account”, and make sure the option to authenticate via Filemaker File is confirmed (**Figure 6C**) (note that the “Account Name” must match the name of the laboratory within FishNet). Type a password for the new account, then select “Privilege Set” and choose “New Privilege Set”. A popup window will open, and within that window change the privilege set name to the name of the laboratory. After the name is changed, make sure all the data access and design options are set to “All modifiable”, then click Records and select “Custom privileges” (**Figure 6D**). A new window will be opened where you can see all of the tables that store information in the database **(Figure 6E)**. Each table will have a different setting. Here, you can select “Fish Lines” then change the “Delete” option to “no”. Select “Tanks” and change the “View” option to “limited” then enter the following text into the popup window, **Laboratories::Lab Name = “Wythe”**, replacing “Wythe” with the desired laboratory. Repeat this same operation for the “Delete” option **(Figure 6E)**. In order to complete the set up for each new laboratory, the administrator must set up the appropriate permissions for all data tables. For a fully functioning lab, refer to **Figure 6D** and modify each field as shown. Follow the previous instructions for every instance of “limited” privileges, referring to **Table 6** for as necessary (**Supplemental Video 4**).

**Table 5.** Hardware, software and cost comparisons between FishNET configurations.

| Application | Numbers of Users Supported | Hardware Requirements | Platform | Contract Length | Cost** |
| --- | --- | --- | --- | --- | --- |
| **FileMaker Pro Advanced (1 instance)** | 1 Computer | For computer running Windows: CPU: 1 GHz or faster x86- or x64-bit processor<br>RAM: 1 GB<br>For computers running macOS: RAM: 2 GB | Windows 10, 9, 8, 7 with latest updates.<br>macOS High Sierra 10.13 or Sierra 10.12 | Single permanent license | \$ 324 |
| **FileMaker Pro Advanced (for teams of 5 or more)** | Allows for 5 simultaneous client connections to one or multiple databases | Same as above | Same as above | Billed annually or permanent license | \$ 540 if billed annually<br>\$1,620 for 5 permanent licenses |
| **FileMaker Server 17** | Allows up to 500* users | CPU: Dual Core<br>RAM: 8 GB<br>Hard Drive: 80 GB or more | Windows Server 2016 Standard Edition (with Desktop Experience),<br>Windows Server 2012 R2 with Update,<br>macOS High Sierra 10.13 or macOS Sierra 10.12. | Included with FMP Advanced Team License | Variable |
| **FileMaker Cloud 1.17** | Allows up to 100* users | NA | NA | Billed annually | \$ 400 (user licenses not included) |
| **FileMaker Go** | NA | iPad or iPhone | iOS 11.2 or above | NA | Free (Apple App Store) |

\*A user license is required to access custom applications hosted by FileMaker Cloud or FileMaker Server from FileMaker Go. The number of FileMaker Go connections may be limited by hardware, app design, operating system, or license agreement.
\*\*Education and non-profit prices shown.

**Table 6.** Setting up lab privileges.

| Table Name | Limited Fields | Calculation |
| --- | --- | --- |
| Tanks | View | Laboratories::Lab Name = “Lab Name” |
|  | Delete | Laboratories::Lab Name = “Lab Name” |
| Fish Crosses | View | Laboratories Crosses::Lab Name = “Lab Name” |
|  | Delete | Laboratories Crosses::Lab Name = “Lab Name” |
| Harvest Assets | View | Laboratories Harvest::Lab Name = "Lab Name" |
|  | Delete | Laboratories Harvest::Lab Name = "Lab Name" |
| Harvests | View | Laboratories Harvest::Lab Name = "Lab Name" |
|  | Delete | Laboratories Harvest::Lab Name = "Lab Name" |
| Lab Members | View | Lab Name = "Lab Name" |
|  | Delete | Lab Name = "Lab Name" |
| Laboratories | View | Lab Name = "Lab Name" |
| Dead Fish | View | Laboratories::Lab Name = "Lab Name" |
| Event | View | Lab = "Lab Name" |
|  | Delete | Lab = "Lab Name" |
| Nursery | View | Laboratories Crosses::Lab Name = "Lab Name" |
|  | Delete | Laboratories Crosses::Lab Name = "Lab Name" |

### Entering Lines

Lines, in this case, refers to distinct alleles that follow ZFIN established nomenclature (https://wiki.zfin.org/display/general/ZFIN+Zebrafish+Nomenclature+Conventions). Novel lines, such as those generated by CRISPR/Cas9 mutagenesis, can be also be entered in FishNET. To enter an allele, from the landing page, select the “Lines” tab (or select the “Fish Lines” layout from the Layout pulldown menu) (**Figure 7A**). If your database has no lines entered, you will be taken to a blank record (**Figure 7B**). Under “Record View” you will only be able to view one line or record (e.g. a unique allele) at a time. Use the forward and backward icons at the top, upper left of the database window to scroll through the records manually (**Figure 7B**).

**Figure 7.**
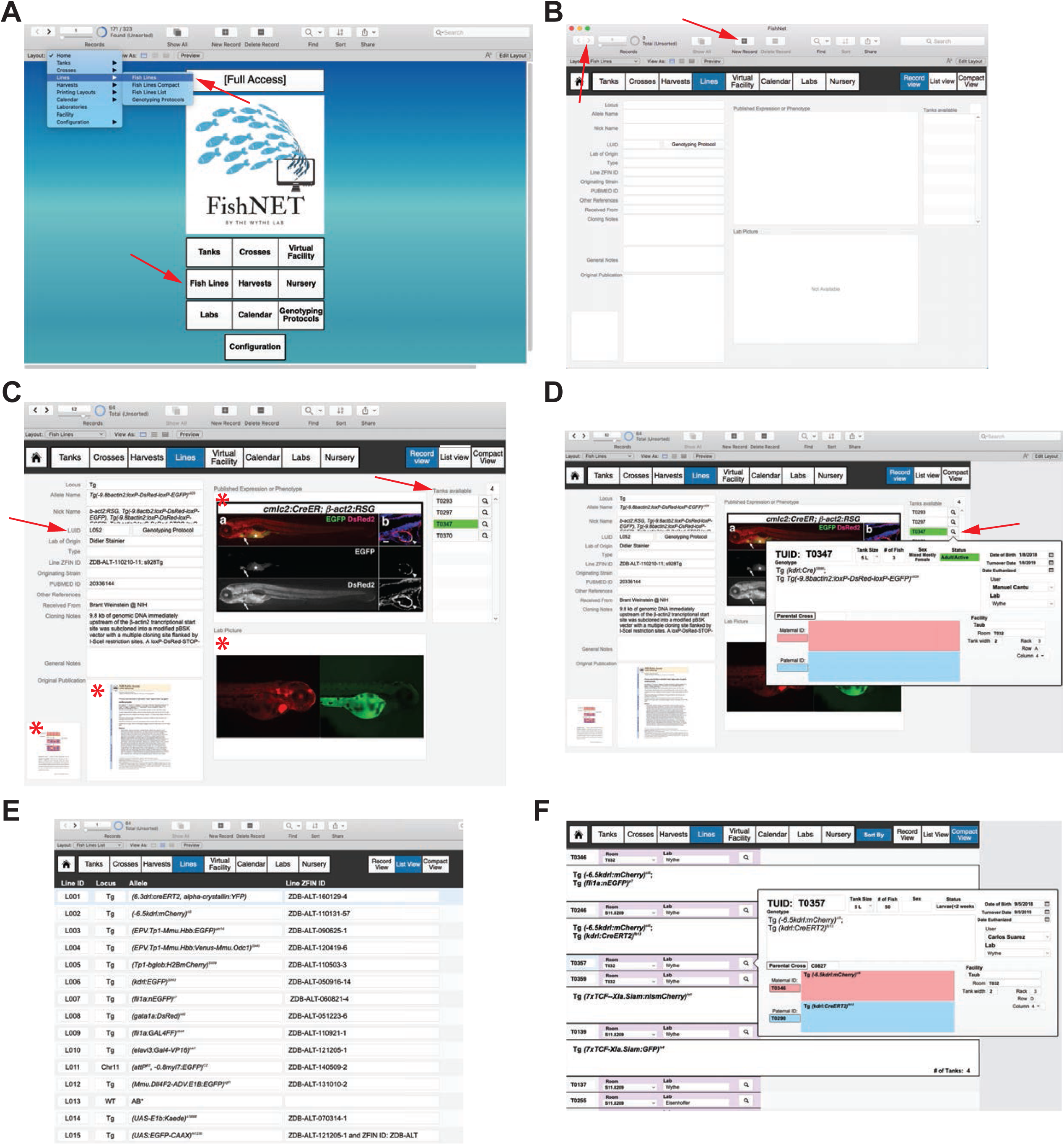
Entering lines. A) The “Fish Lines” layout can be accessed from main layout drop-down menu, or by clicking “Fish Lines” on the main landing page (each indicated by the red arrows). B) A new line information page can be created by selecting “New Record” (indicated by the red arrow in the header). Users can move through all records using the forward and backward arrows (top left of the header, indicated by the red arrow). C) Each new record is assigned a Line Unique ID (LUID), indicated by the red arrow. This is an example of a fully annotated line record. Containers where external pictures or file attachments can be placed are indicated with an asterisk. Any tanks available for a given line will be displayed on the far-right side of the main window under the “Tanks Available” label. The tank labels are color coded according to their age and mating status. D) Selecting the magnifying glass will bring up a popup window with a detailed overview of an individual tank. Lines can also be browsed in list view (E) or compact view (F) for easy navigation.

To create an allele in “Record View”, populate the following fields in the blank record: locus, allele name, nick name, line ID, lab of origin, line ZFIN ID, originating strain, PUBMED ID, laboratory the line was received from, cloning notes, and general notes (**Figure 7C**). Importantly, whatever designation is used for the “Allele Name” field will be redeployed throughout the database in dropdown menu choices, ensuring uniform nomenclature within a research groups, across other labs, and throughout entire facility (**Supplemental video 5**). Each line that is entered is assigned a Line Unique ID number (beginning with L). Each subsequent record is automatically assigned the next available Line UID number (L+1) (**Figure 7C**). FishNET also allows for users to upload and store attachments associated with each line, such as associated publications, representative images of example phenotypes or expression patterns that can function as a reference for sorting offspring (**Figure 7C,** indicated by asterisks). Additionally, one can link genotyping protocols to individual lines, such that the protocol can be accessed by selecting the “Genotyping Protocols” layout or button within the “Record View” field (these will be discussed in more detail in the following section).

While users or owners of the line are not defined when a line is created, later, if tanks are populated with a line and placed on a rack, they will be visible in the “Lines: Record View” window under the “Tanks Available” heading. Only live tanks will appear within this window. Clicking on the magnifying glass provides a detailed view of the tank, including its genotype, date of birth, sex, number, the owner of the line, its status (e.g. juvenile, adult), location in the facility, and the paternal and maternal animals that generated the tank (**Figure 7D**).

To view all lines that have been entered into a database, within the “Lines” layout select “List View” (rather than the individual “Record View”). This generates a simple, scrollable list of every allele, including the locus, allele, and ZFIN ID. To define a new locus, simply input text into the space provided, and the new locus will be indexed and will appear as an option in any subsequent input. To enter a new line, type in the correct genetic manipulation under the allele field, and the rest of the database will populate the allele name using this designation. Within “List View”, users can enter in a new line by selecting “New Record” and entering in the Locus, Allele, and ZFIN information. For adding additional details, select the “Record View” layout.

To view all lines that are actually available within a facility or user group, select “Compact View”. Users can navigate this view using the “Sort By” button to segregate the animals by any of the following criteria: genotype, facility, lab, or user. This view will show users the TUID, Facility, and Lab that own any tank(s) of a given genotype. Additionally, users can click on the magnifying glass for a detailed view of that particular tank (location, parents, age, etc.) (**Figure 7E**).

### Entering Genotyping Protocols

To create a genotyping protocol, from the landing page select “Genotyping Protocols” (**Figure 8A**), then select the “New Record” button. Here you will name the protocol, enter the ZFIN identification number and any notes, as well as PCR primers and thermocycler conditions, expected product size, and a representative agarose gel image by dragging and dropping any .PNG or .JPEG file into the PCR example container (**Figure 8B**). For optimal viewing, gel images should be resized to 4 inches width and a resolution of 300 pixels/inch. To scroll through available protocols, use the FileMaker forward and backward keys available at the top of the screen, or use the search window.

**Figure 8.**
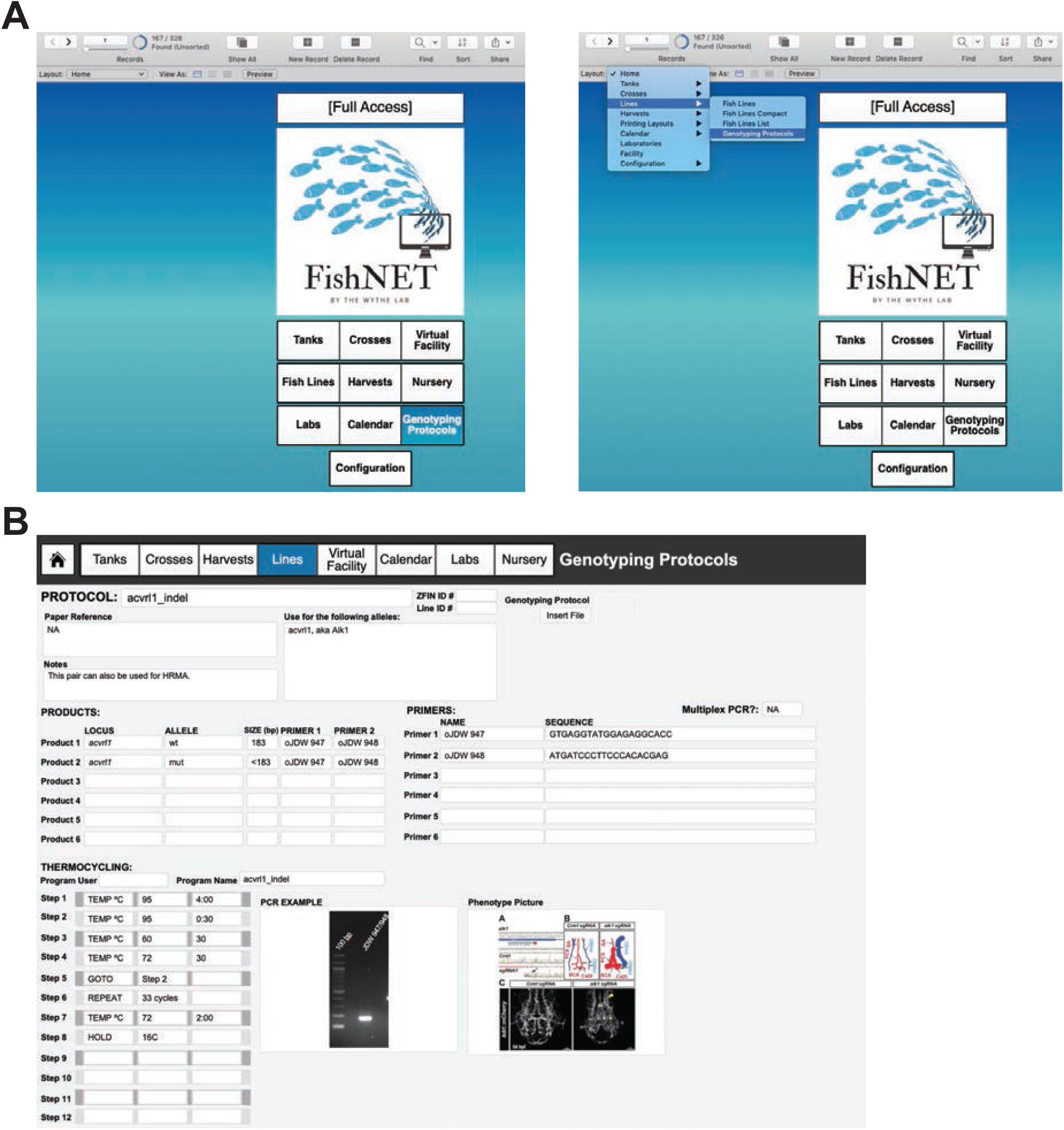
Entering genotyping protocols. A) Genotyping protocols can be accessed from the landing page via the “Genotyping Protocols button (left side), or via the dropdown menu under “Lines” and “Genotyping Protocols” (right side). C) An example genotyping protocol, where primers, PCR conditions and images, as well as phenotypic information can be imported.

### Creating, Locating, Moving, Removing, and Retiring Tanks

The underlying logic of FishNET that enables the tracking and searchable features relies upon a Tank Unique Identifier (TUID) that is assigned to every new tank. From the “Tank List” layout (Tanks>List View or Landing Page>Tanks) one can see a list of all Tanks, each with a TUID, stored within the database (**Figure 9A, B**). All pertinent information (sex, genotype, age, owner and location) are present in this view. To create a tank, click the “Detailed View” tab (or from the main dropdown menu select “Tanks” and go to “Tanks Detailed View”). In that layout, select “New Record”. Then select the loci, and the appropriate allele, from the drop-down list (that should have been populated by following the instructions outlined in <u>Entering Lines</u>). After this, select the “User” of the tank (this will auto populate the user’s lab, email address, institution, phone number, and IACUC information). If the TUID of the maternal or paternal parent tank is known, enter it now to establish a pedigree for this new tank. The TUIDs can be entered manually, or you can select the field(s) with your cursor and then scan a barcode for the animal of interest. Be sure to enter the date of birth, tank size, and # of fish for the TUID record. The status of the line will be color coded according to the date of birth (fish less than 3 months remain white; fish older than one month, but younger than one year, are green; fish older than 1 year are red; and retired or euthanized tanks are black). Additionally, the database automatically creates a turnover date for one year after the date of birth. Users can choose to receive an automated email reminder for line turnover by selecting the “Set reminder” calendar button above the “Turnover Date” field (but this must be done after establishing the User/Owner of the tank to set up an email address for the reminder). Finally, a user must assign a location for the tank by selecting the “Facility” drop-down menu and entering in the Rack, Row, and Column for the new tank. After entering that information, select the “Add to Rack” button to place the tank in the virtual facility. Alternatively, one can click the “Add to Rack” button in the header field and then scan in the barcode location on the rack followed by the barcode on the tank to establish the location of the tank (**Supplemental video 6**). Once the user adds the tank to a rack, selecting the Locate Tank button will show the virtual rack window where the tank in question is located (**Figure 9C**).

**Figure 9.**
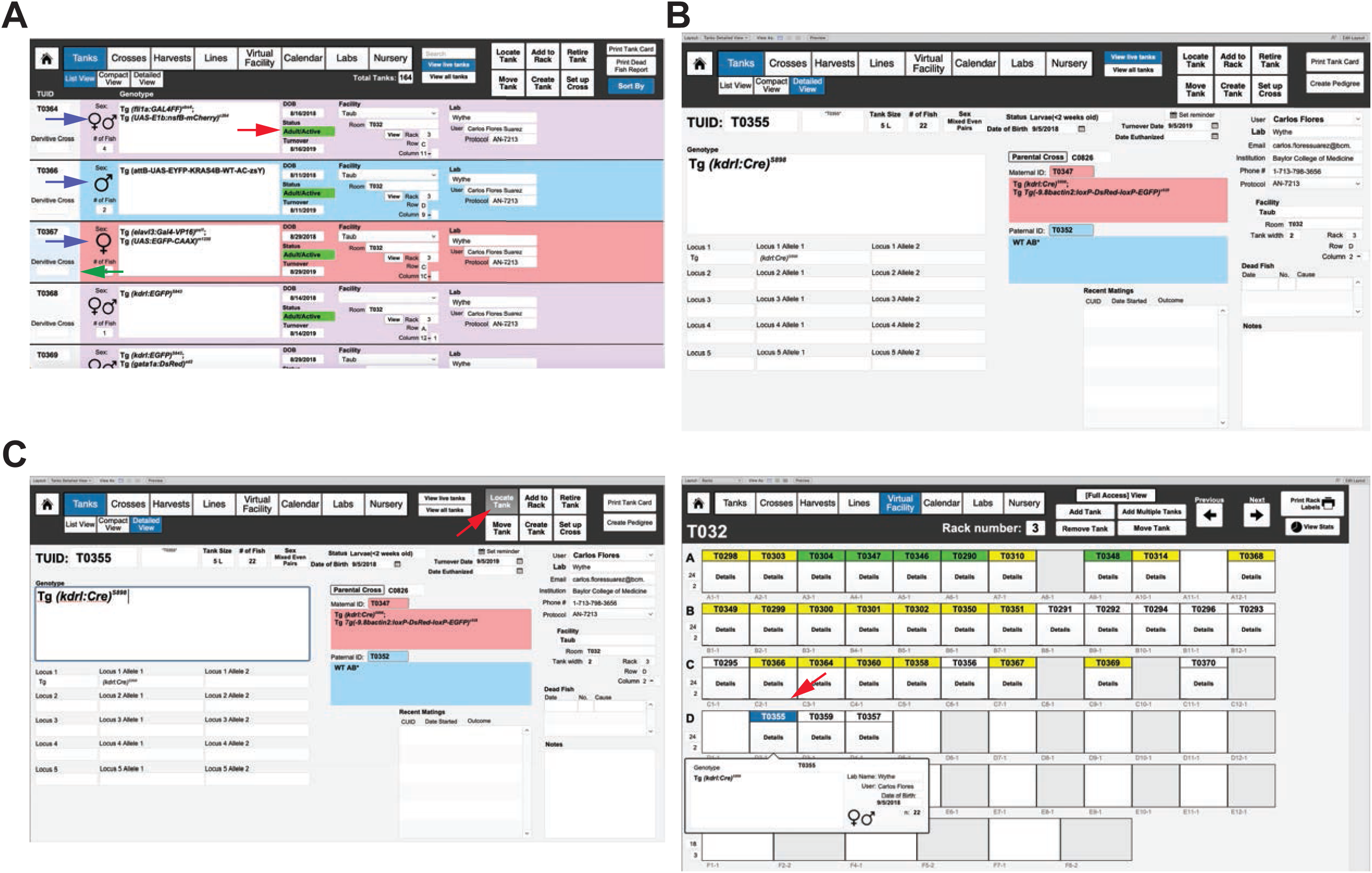
Entering and viewing tanks. A) **“**List view” of all “live” (active) tanks, in this case sorted by “Facility”. All active tabs for viewing are indicated by blue coloring. The age and status of each tank is indicated by color (see red arrow), while the sex of each tank is indicated by the background color (blue for male, pink for female, and purple/pink for mixed sex tanks, indicated with blue arrows). Selected tank is indicated by a bar changed to light blue (indicated by green arrow). B) Detailed view of a tank with the derivative cross (C0826) and genotype/TUID of the parents, date of birth, date for turnover, and user/owner of the tank all indicated, as well as the facility and position. C) To view the virtual position of the tank, select “Locate Tank” (indicated by the red arrow) in the “Tanks Detailed View” layout, which will bring you to a new window (on the right). The Room will be indicated on the upper left of the header menu (in this case, T032), while the Rack will be indicated in the center of the header menu (in this case, rack 3). The position of the tank of interest will be indicated by a blue label. In this case, TUID 0335 is in Room T32, Rack 3, Row D, position D2-1. Clicking on “Details” will show the genotype, sex, age, and number of fish in the tank, as well as the owner of the tank. All other tanks are colored according to their age and mating status (white = < 2 weeks old, yellow = < 3 months old/juvenile, green = > 3 months old/active, and red = > 1 year old).

### Setting up Crosses

Each mating generates a Cross Unique Identifier number (CUID) to track every mating and to generate pedigrees. To create a cross, select the “Crosses” button from the header or landing page menu (or select “Cross” from the pulldown menu). Then, in the header menu, select “Set up Cross”. A pop-up window will appear asking for the paternal TUID and the maternal TUID (**Figure 10A-C**). These can be entered either manually by text, or by reading the barcode of each parent tank. Next, a pop-up window will ask what type of mating is being setting up: a bulk cross, a trio mating (defined as one male crossed to two females), or a pair. The next window will ask how many of this type of mating are being setting up (e.g. 8 pairs). After filling in this field, the application will show you a mating label, while asking you if you want to repeat the cross. If you are only setting up one type of cross (e.g. just pairs) and selected the appropriate number, then select “No”. If you wish to set up another type of mating from this cross, such as a few trio tanks, then select “yes”. The cycle repeats until you chose to no longer repeat this cross. After choosing this option, the screen will return you to the “Crosses” layout. To simplify the view, you can choose to “view active crosses” rather than “view all crosses”. In the header field you will now have the option to print mating labels for each of the individual mating tanks that you set up. These labels display the genotype, sex, TUID, and location of the parent tanks, as well as the owner of the cross. If additional mating labels are needed, select the “Print Additional Labels” field in the header. We also use these to label 10 cm plates of embryos from crosses. Active crosses that are less than 1 day old will be labelled green, while active crosses more than 1 day old will be labelled red to indicate that the fish should be returned to the system (or fed). Retired crosses will be colored grey (and every cross should be retired once the fish are taken down and returned to the system). This convenient color-coding system enables easy visual tracking of active crosses within a facility or user group (**Figure 10D**) (**Supplemental video 7**).

**Figure 10.**
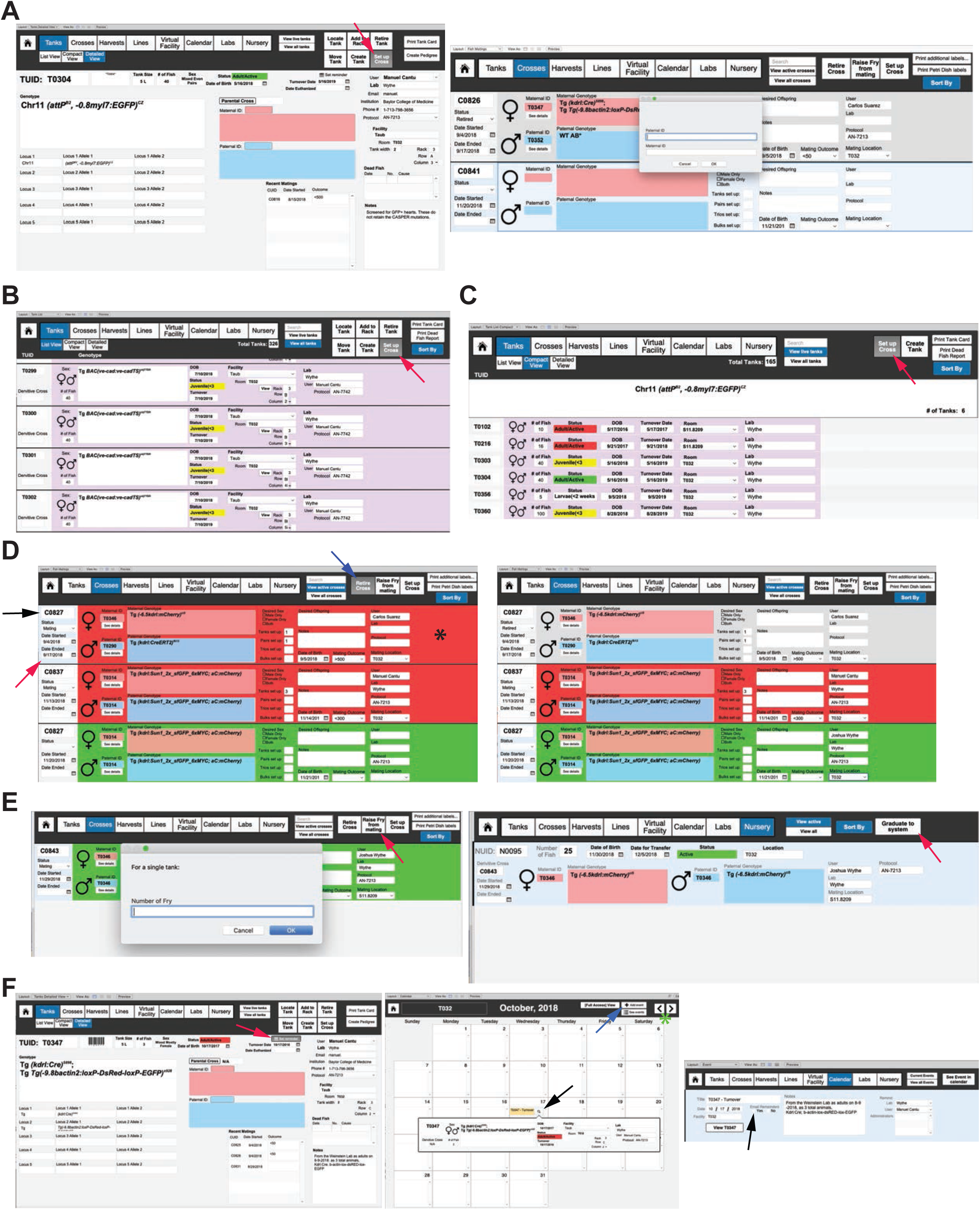
Setting up crosses and calendar reminders. A) Crosses can be set up from the “Detailed View” by selecting “Set up Cross” button (far left screen image), indicated by the red arrow, which will take you to a new screen (far right screen image) where you can either manually, or by barcode reader, enter the TUID for the dam and the sire for the cross. Selecting the “Set up Cross” function automatically generates a new Cross Unique ID number (CUID) and record for tracking purposes. The crossing function also allows you to record the success of the mating for tracking the fecundity of a tank (and overall mating at the level of a rack, room, or facility) (not shown). B-C) Crosses can also be set up in the “Tanks List View” layout (B) or in the “Tanks Compact View” display (C), which will take the user to the same popup window asking for maternal and paternal TUIDs for the new cross (as in (A)). D) The “Crosses” layout displays either “active” or “all” crosses. The far-left column (indicated by the black arrow) displays the unique cross identifier (CUID) of a cross at the top, with the status of the cross (either mating or retired) indicated below in the dropdown menu. Below that are the dates the mating was started, as well as the date the mating was ended. The blue background in the far-left column (indicated by the far left, red arrow) shows users which cross is currently selected (compare the blue background indicated by the red arrow in C0827 to the light grey background in the far left of C0837 for reference). Each cross is also color coded for easy visual reference (indicated by the asterisk on the far-right side). Green = an active cross, red = an active cross, but more than 2 days have passed since it was set up, and gray = retired crosses. After a cross has been ended, in the top right corner a user then selects to retire a cross (indicated by blue arrow), after which the background color on of a cross will automatically change to grey (compare the background color of C08277 between the left and the right images). The upper right header menu also has options to raise fry from the mating, to set up a new cross, to print more cross labels, or to print labels for petri dishes for embryos. E) Selecting “Raise Fry from Mating” (indicated by the red arrow) will ask the user for the number of embryos that will be kept in each tank/petri dish for raising. After selecting “OK”, a second window will ask if there are additional embryos to raise from the same cross (not shown). Once the user indicates that no more fry will be raised, a new nursing unique identifier record (right side image) will be created. Here, the nursery unique identified record (NUID), as well as the identity of the sire and dam, and number of fish, and other details will be auto populated from the CUID record. Once the fry are ready to move onto the main system, the user can select “Graduate to system” (indicated by the red arrow in the far right image). This function creates a new Tank Unique ID (TUID) autopopulated with all of the pertinent NUID and CUID information (not shown). F) Once a new TUID is created, a user can automatically set a reminder to turn the line over one year from the date of birth by selecting the “set reminder” button (indicated by the red arrow in the far left image). The middle image shows the “Calendar” layout (in this case for the month of October, 2018 for room T032). More “events” can be added (using the button indicated by the blue arrow). Clicking on the magnifying glass (denoted by the black arrow) will create up a popup window showing the details of an event. Months can be changed using the forward and backward buttons on the far right of the top header menu (indicated by the green asterisk). The far right image shows the detailed “event” layout, where FishNET has the capacity to email a reminder to the lab user and/or room administrator for any given event (using the contact details supplied in the laboratory set up).

### Raising Offspring/Graduating Fish and Nursery

FishNET also has the capacity to track all larvae that are destined to be graduated to the main system, enabling real-time tracking of all tanks on a nursery, as well as the success of fish survival rates through the “Statistics” button within the “Virtual Facility” layout. To raise fry from a cross, select the “Raise Fry from mating” button in the header menu (**Figure 10E**). A popup window will ask you for the date of birth for the fry, as well as the number of fry for a single tank to raise (this will help determine survival on the system, as discussed later). Within the “Nursery” layout, each tank of fry is assigned a Nursery Unique Identifier (NUID). After filling in this information, a new popup window will ask if you wish to raise more fry from this cross (e.g. additional plates/tanks).

At day 5, select the “Graduate to System” button in the top right of the header. Upon selecting this button (located in the header field of the “Nursery” layout), a popout window will ask the user to enter NUID(s) that will be used to form a new Tank (in the event that more than one dish of fry of the same genotype are used to create a new tank on the system) (**Figure 10E, right**). This automatically generates a new, unique TUID record that contains all of the previous NUID information (allele, owner, strain, etc.). Once a new TUID is created, a user may select the “Add to Rack” function, as described previously, to “place” the tank to the virtual main system (**Supplemental video 8**).

### Calendar

The calendar layout/function (**Figure 10F**) enables users to access a Facility wide calendar with all related events (crosses set up, system reminders such as change baffle size, alternate food, graduate to main system, etc.). Selecting “Add Event” will take the user to a field where the title of the event, date, facility are entered. Finally, the calendar function has an option for email reminders that automatically populates information using the laboratory user field(s). This function can also be used to copy administrators or animal husbandry staff on reminders.

### Harvests

Another novel feature of FishNET is the ability to track the pedigree of embryos, or adults, that are used for experiments (as well as all experimental conditions) in the “Harvests” layout. This layout generates a novel Harvest Unique Identifier (HUID) that pulls from either the parental cross CUID (for embryos) or the TUID (for an adult) to create a unique record with the date of the harvest, experimental treatment, and number of fish collected **(Figure 11A)**. Additionally, using the “Create Resources” tab, one can generate detailed records for every embryo or adult within a harvest (**Figure 11B**). While non-essential for facility management, many individual labs may find this functionality helpful for tracking embryos or adults that have been harvested and stored in the freezer, keeping track of sections, mRNA, or protein samples. Tagging samples with a unique Resource ID and labelling a sample “H0001-1” is easier and clearer than writing out an entire genotype, sex, date and treatment on one slide, or an Eppendorf tube. Furthermore, there will be a permanent record of the parents, animal, genotype, and treatment associated with this HUID. Additionally, images (such as genotyping gels) and other relevant data can also be pasted within these “resource” pages (**Figure 11C**).

**Figure 11.**
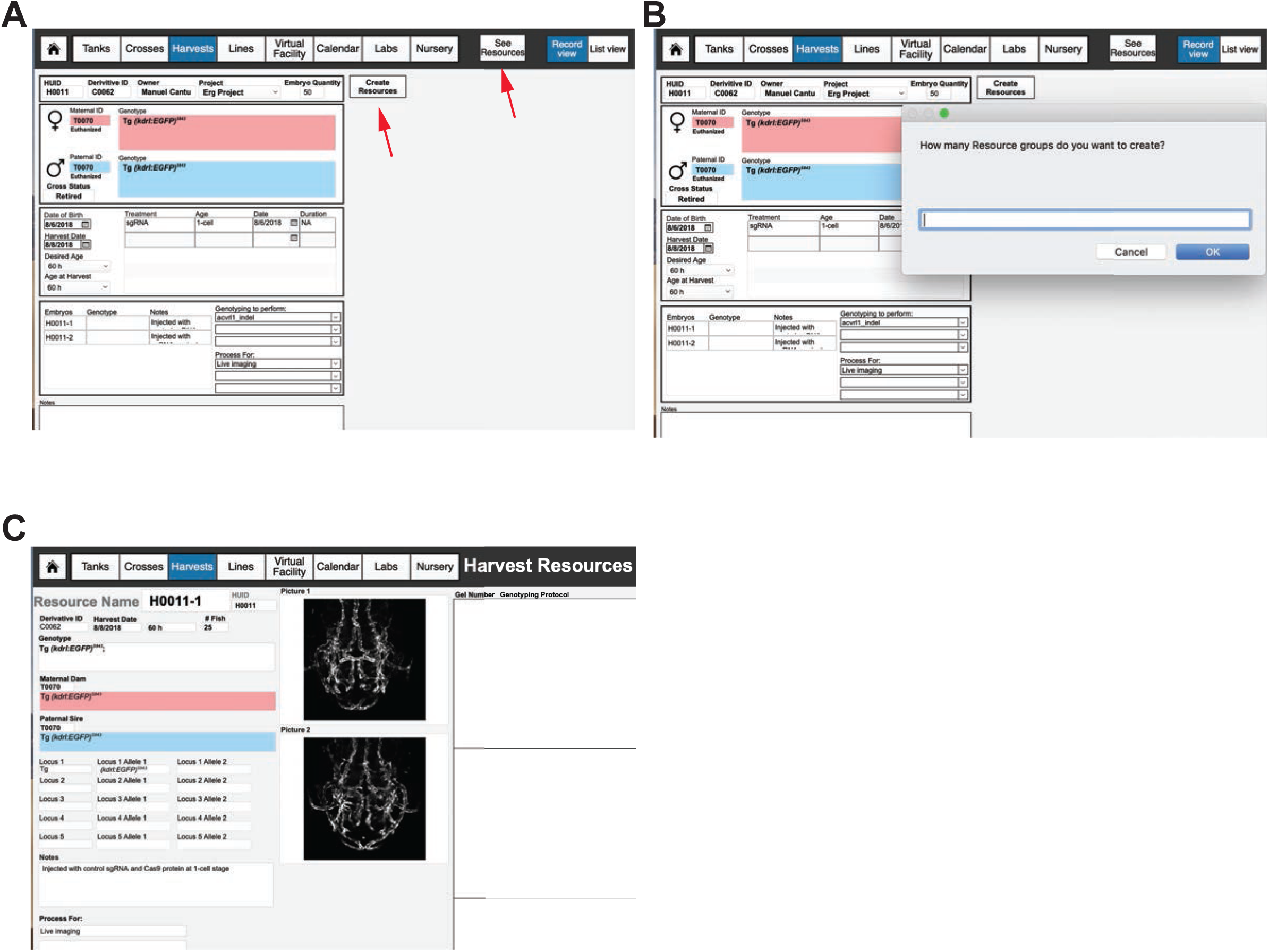
Using FishNET to track harvest records. A) Embryo harvest template. Creating a new record in the “Harvests” layout will create a new Harvest Unique ID record (HUID), where information such as the cross used to produce embryos, the genotypes and TUIDs of the parents, the date of collection, and number of embryos can be entered. Information regarding any chemical or genetic treatments, the age of collection, any genotyping to perform, and downstream tissue processing can be entered into this record as well. B) After an embryo/adult quantity is added, a user can select the “Create Resources” tab to create a unique Harvest ID (H000X-1 if one resource, 1 and 2 if two, and so on) for each embryo group or adult fish that is processed. Selecting “Create Resources” will prompt the user to select the number of groups that embryos or adults are going to be divided into. If the original embryo quantity is 50, the user could make 2 groups of 25 embryos each with a different experimental condition. C) Viewing the resources takes the user to an individual record view of all “resources” associated with that harvest. Information regarding genotype, tissue processing and images of the embryos can store here to create permanent records of any experimental harvest.

### Automated Genotype Annotation for Harvest

To automate annotation of genotyping gels, we created an Apple script (FishNET_Genotype.scpt file) that can take the record list from the harvest section of FileMaker Pro Advanced into Adobe Photoshop Creative Cloud 2017 to annotate images of PCR gels (this function is only available in Mac OS X). This requires a completed genotyping protocol (**Supplemental Figure 1A**), which can be found under Lines/Genotyping Protocols. A special security permission is also needed to allow Apple scripts in FileMaker Pro Advanced, Select File / Manage / Security… Select Extended Privileges and give full access to “fmextscriptaccess”. To start, download the script (**Supplemental File 1**). Open the script in Script Editor and follow the simple instructions to adapt it to your computer (**Supplemental Figure 1B**). We recommended adding a shortcut for the script to your menu bar. From the script button on the menu bar, select “Open Scripts Folder / Open Users Script Folder” and deposit the downloaded script file there (now the script can be run from the Script menu in the menu bar). Before running the script, a genotyping gel should be open and resized in Photoshop to 4 inches width by 3 inches wide, with a resolution of 300 pixels/inch (**Supplemental Figure 1C**). Run the script and enter the Harvest Unique ID number you want to annotate. Make sure all names match the correct bands on the image, then select “OK” in the pop-up window asking “Are ready to add the gel to the database?”. Select “Yes” to create a new genotyping record. Drop the new annotated gel image into the empty field and select the appropriate name of the genotyping protocol as seen in **Supplemental Figure 1D**. The annotated image is now present in the detailed view of all records linked to the selected harvest ID (**Supplemental Figure 1E**).

### Importing Existing Data to FishNET from Excel or FileMaker

As many laboratory groups use excel spreadsheets to manage their existing colony, we provide a simple way to import this information into FishNET. Once the laboratory, members, facility and rooms are set up in FishNET, one can import preexisting line data. In excel, ensure that the following columns are populated: Locus (Tg, Et, etc), Allele Name, Lab of Origin, and General Notes (**Figure 12A** and **Supplemental File 1**). After all fields are entered, save the document in Tab delimited text (txt) format. To import these data into FileMaker, go to the “Lines” tab, select File/Import Records/ File. Next, select the .txt file where the data was saved. Once selected, the user has to match the source field from excel to the target fields in FileMaker Pro. Activate field mapping and importing by selecting all fields that should be imported (the middle symbol between “Source Fields” and “Target Fields” should change to an arrow) (**Figure 12B**). “Add new records” should be selected, as well as “Don’t import first record (contains field names)” (**Figure 12B**). Similarly, it is possible to import data from a pre-existing FileMaker database. In the original database, go to the layout you wish to export (Tank list, Lines, etc) and select File/Export Records. Then, “save” the new file as a FileMaker Pro Advanced Type (fmp12) file. Next, select the fields to be exported and select “Export”. Finally, to import these data into FishNet, follow the above instructions selecting the new .fmp12 file as the source of import.

**Figure 12.**
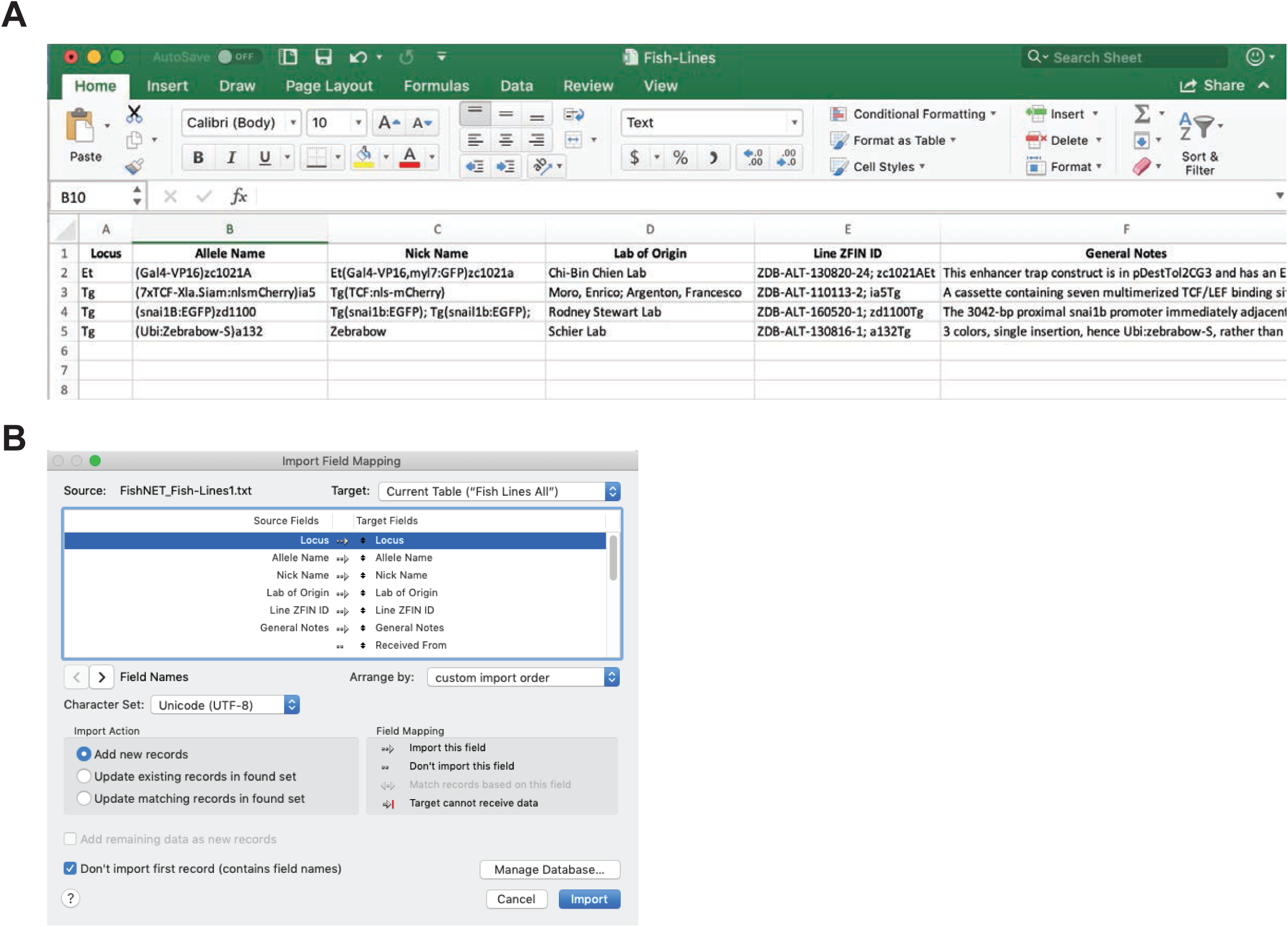
Importing data into FishNET. A) Screenshot of the excel template used to import lines into FishNET. All fields should be filled accordingly. B) Import Field Mapping showing the correct configuration to import new lines into FishNET.

### Reporting Capabilities

We placed an emphasis on the reporting capabilities of the database, which allow for storing fish census and usage reports for the whole facility in an exportable, excel-compatible format file. This graphical report provides an overview of all fish morbidities and mortalities, and individual fecundity records are maintained for each line across the entire facility. FishNET also tracks all animals, can generate user defined fish use reports (as required for annual IACUC protocol reporting), and as previously mentioned generates graphical reports for water quality data. Within the “Virtual Facility” layout, selecting the “View Stats” function **(Figure 13A)** will take users to a graphical report of the number of tanks and their status (i.e. adult mating age, juvenile, older than one year) and the number of total fish and tanks (**Figure 13B**). Due to limitations in graphing capabilities within FMPA, we have opted to generate reports using Google Charts Application Programming Interface (API), a third-party software (as outlined below in the Importing Water Quality section). Selecting the “Vitality Summary” button in this layout will display the number of births, natural deaths, and euthanized fish across a rack, room, or facility with weekly, monthly or yearly reports (**Figure 13C**). These graphical reports for facility stats and fish stats are both populated by user entered data from dead fish reports, mating records, and tanks that are retired. Finally, in the individual room view, there is also a “Water Quality” report that will generate graphical views of user-uploaded water quality reports detailing the conductivity, pH and temperature for the racks within a room (**Figure 13D**, described in detail below). In this “View Stats” layout, as different systems have unique headers and labels for data, we have simplified uploading external data. The management and partitioning of users and labs in FishNET, combined with barcode labeling, will allow the PIs and animal facility managers (or staff) to track animal usage and husbandry and enable accurate billing and IACUC reporting in real-time. FishNET also has the ability to create a weekly report of fish found dead in the system. In the tanks template, select “Print Dead Fish Report” (**Figure 13E**). Select the desired week to create the report and facility and print (**Figure 13F**).

**Figure 13.**
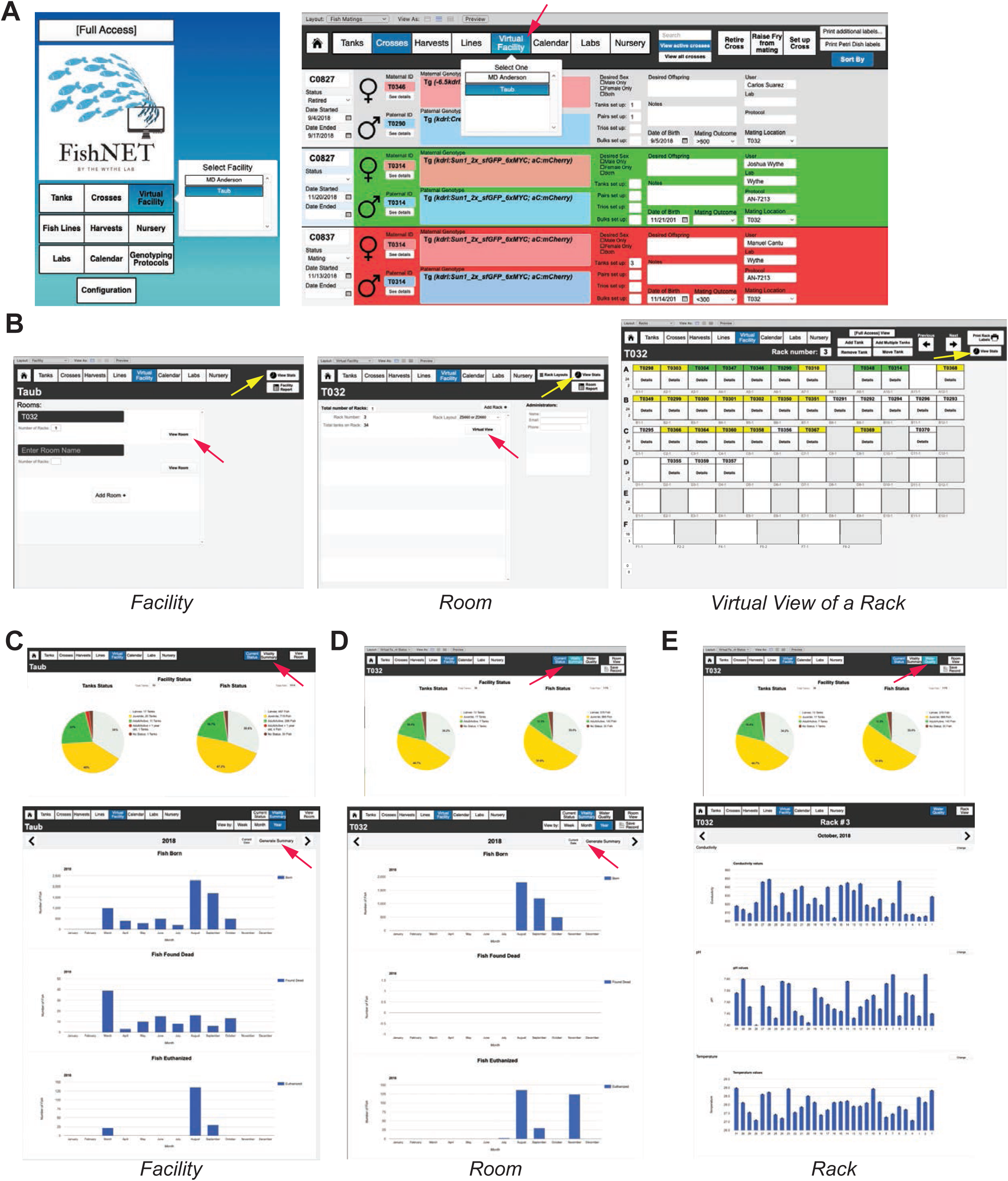
Generating reports with FishNET. A) Navigate to the “Virtual Facility” from either the landing page (left) or the header tabs ^12^. B) From the “Facility” page, choose a “Room”, then select “Virtual View” to visualize the rack and all tanks within the rack (far left). Selecting “View Stats” (yellow arrows) in any of these three windows will take the user to a statistical report for either the facility, room, or individual rack. C) Shows the “View Stats” layout for an entire facility, with the “Current Status” of all tanks in that facility on the left, and all individual fish on the right. From here, one can also view the “Vitality Summary” (indicated by the red arrow in the header menu) for the entire facility, room, or rack. D) “Generate Summary” (indicated by the red arrow) in the “Vitality Facility Summary” layout creates a report either for the past year, month, or week displaying all recorded fish born (embryos) from crosses, dead fish found in the facility, and fish euthanized, for a comprehensive view of the vitality of a facility or room. E) Water quality graphs showing conductivity, pH and temperature of individual racks. Water quality can also display an average value of all racks in one room.

### Importing Water Quality Records

We have enabled FishNet to generate water quality reports (e.g. conductance, pH, etc.) using the interactive Web service, Google Charts. Due to the limited options for data display in FileMaker, we have instead used Google Charts to generate graphical reports, as it is more flexible in terms of graphing options, colors, and data input options. To generate these visual reports, export existing system water quality data as a .CSV file from a rack or facility of interest. Then, open FishNET and navigate to “Virtual Facility” and the specific “Room” containing the rack that the data corresponds to. Next, select “Virtual View” for the individual rack of interest, then in the next window select “View Stats” (**Figure 14A**). Once in the individual rack stats view, and in the present month of interest for the data, select “Water Quality Report” and “create new record” to import data (**Figure 14B**). Two different fields with either “Days” or “Values” will appear for each parameter field (conductivity, pH and temperature). From the .CSV file, simply copy the date of data acquisition (dd/mm/yy, or just the day number) in the first field and enter the values in the second field (making sure that the number of days and values are correctly related to one another). If there are multiple values recorded on one day, the system will display a bar graph with the average values and corresponding error bars.

**Figure 14.**
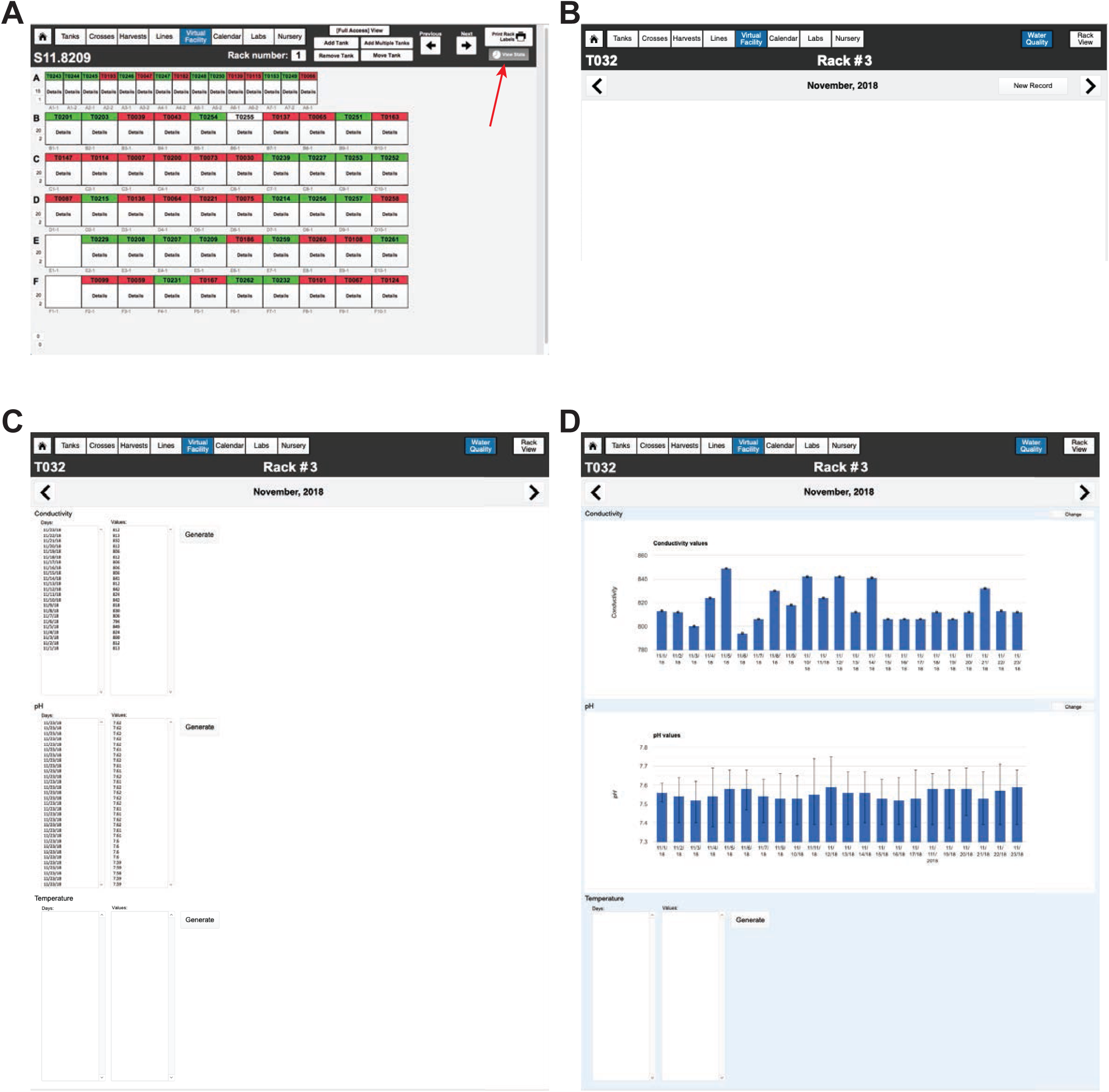
Importing water quality records. A) Example of virtual rack view, to see rack statistics, select View Stats (shown with red arrow) on the top right corner of the screen. B) Scroll through the months with the arrows in each side of the header. Once in the current month, select “New Record”. C) Days and values can be pasted in the two fields, FishNET can show either a single value per day (conductivity) or the average of multiple readings a day ^31^. D) FishNET uses Google Charts to display data and for that reason an internet connection is needed to use the graphing functions. Once the user is done adding the days and values, select “Generate”. Graphs can be modified by selecting “Change”.

### Printing Tank, Nursery, and Rack Labels

During the course of optimizing FishNET we tested numerous printer configurations to find a sturdy, waterproof re-adhesive label that is also barcode compatible in a size compatible with large and small aquarium tanks and 10 cm petri dishes for fry. After much trial and error, we recommend using a BBP33 Brady printer and permanent, bar-coded polyester labels for each tank location label on a rack. Unfortunately, the BBP33 printer only supports Windows computers. While some third-party vendors have developed BBP33 drivers for Mac, those we tested were unreliable. Instead, as a workaround we suggest installing Windows 10 using Parallels Desktop for Mac (https://www.parallels.com/). As a cheaper alternative, one may use a Dymo 450 LabelWriter Turbo printer, but note that the labels are permanent, and their removal may require scratching the tanks (details available, as well as printer layouts, upon request). Printing layouts for various labels are provided within FishNET under the main layout dropdown menu under the heading “Printing Layouts”. From here, users can visualize all of the printer layouts for petri dishes (for embryos), fry on the nursery, small tanks, large tanks, racks, and crosses. Each of these layouts are modifiable and the label size can be changed to one that best meets a user’s needs, including which information should or should not be present on the labels. To create barcodes in FishNET, the Code 39 Barcode Font from IDAutomation (IDAutomationHC39M Code 39 Barcode.ttf) is required in any computer printing barcodes. It can be downloaded for free here: https://www.dafont.com/idautomationhc39m.font. Once the font is properly installed, the user needs to go to each printing layout (**Supplemental Figure 2A**) and select “Edit Layout”. In edit mode, select the barcode rectangle and change its font from Arial to IDAutomationHC39M (**Supplemental Figure 2B**). A barcode should appear. Then “exit” the layout and “save” changes. Repeat this process with each printing template.

After configuring a facility, within the “Virtual Facility” layout a user can select their “Facility”, then select an individual “Rack”, and within this layout a button to “Print Coordinates Labels” is present in the upper right area of the header. Selecting this function will create a pop-up window asking, “Which Row do you want to Print?”. After entering the desired row of the rack you’re currently using (e.g. “A”), then selecting “OK”, the labels for one row will be printed. For labelling the tanks on the main system and nursery, as well as the petri dishes containing embryos, we prefer to use a removable label, which has the added benefit of reducing adhesive build up on tanks, while also eliminating errors in transferring information (such as D.O.B., genotype, sex, etc.) that occur during manual transcribing of labels. For an example of a completely labeled rack and tanks see **Supplemental Figure 2C-D**. To make input of tank locations easy, a barcode compatible function is ready using the “Add Multiple Tanks” function in the Virtual Rack template, where a sequential series of pop up messages will prompt the user to scan the tank barcode followed by the rack location barcode (printed above).

### Frequently Asked Questions

A common error in FMPA (at least within our team) is after importing data, or deleting a record, a user would like to reset the TUID, CUID, LUID, or NUID number to a previous value (to eliminate gaps in numbering and ensure consistency). To do so, select “File” in the Filemaker Pro Advanced menu, go to “Manage” then select “Database”, and a pop-up window will appear. Within this pop-up window, go to the “Tables” section and double click the table you want to edit (e.g. “Fish Crosses”). Then double click the unique identifier Field Name (e.g. “CUID”), and a pop-up window with options for the field will appear. Then, select the “Auto-Enter” section. Here, the “Serial Number” option will have a checkmark, and below this option the next serial value is defined (e.g. “C0032” if you are in “CUID”). To change this next value, simply edit the number after the first character (e.g. change “C0032” to “C0024”). After the changes are made, click the “OK” button to close the “Field Options” pop-up window, then select “OK” again to close the “Manage Database” window, and then finally save the changes.

## DISCUSSION

Despite the increasing demand to ensure rigor and reproducibility at each step of the research endeavor, a robust, affordable, and intuitive archival database for zebrafish animal husbandry records has not been developed and widely adopted by the zebrafish community. We have created a facile, network-accessible relational, open source database that meets the needs of researchers, animal husbandry staff, and institutional animal oversight committee members alike that creates and preserves comprehensive, detailed records for an individual lab or entire zebrafish facility. Additionally, such centralized, comprehensive record keeping, and the data visualization tools contained within FishNET, will limit unnecessary duplication of lines within and across laboratories, ensure timely line turnover, and should flag fish husbandry or facility-wide issues (e.g. water quality, decreased fecundity, etc.) in real-time, allowing for institutions to reduce the overall number of animals required for experiments, and save critical research dollars.

Using the open-source, non-coding-based FMPA platform, the underlying architecture of FishNET can be modified by any end user to meet their unique needs, or directly expanded upon to increase database functionality. FishNET also scales according to user demand, as does the FMPA platform, enabling functionality for research groups as small as one laboratory, or as large as an entire institution. Continual upgrades, along with integration of improved FMPA software and computer hardware technologies will ensure that this inventory system continually evolves to meet the needs of the zebrafish community.

## Supporting information

## ACKNOWLEDGEMENTS

We thank the animal husbandry staff at Baylor College of Medicine, and Philip Kahan in the Eisenhoffer lab, for excellent care and maintenance of our zebrafish colonies.

## AUTHOR CONTRIBUTIONS

Conceptualization: J.D.W.; Methodology: A.C.G., M.C.G., A.M.R., J.D.W.; Investigation: A.C.G., M.C.G., A.M.R., O.E.R., G.T.E., J.D.W.; Resources: A.C.G., M.C.G., J.D.W.; Writing – original draft: A.C.G., M.C.G., J.D.W.; Writing – review & editing: A.C.G., M.C.G., O.E.R., G.T.E., J.D.W.; Visualization: A.C.G., M.C.G., J.D.W.; Supervision: G.T.E, J.D.W; Project administration: M.C.G., J.D.W.; Funding acquisition: M.C.G., G.T.E., J.D.W.

---

**Supplemental Figure 1. Automated genotype annotation.** A) Overview of a genotyping protocol. Expected products, primers and thermocycling conditions, along a PCR example and external document can be imported. B) AppleScript editor settings window with a Script Menu activated. C) Genotyping gel in Adobe Photoshop CC 2017 with the Image Size windows showing the new width (4 in) and resolution (300 pixels/in) needed to proper gen annotation. D) Annotated gel imported in FishNET, the list of resources to which the gel is linked to are listed on the right. E) Example of an individual resource detailed record (H0002-1) with annotated genotype shown on the right.

**Supplemental Figure 2. Printing labels with barcodes.** A) Label for crosses shown, with the layout path to get to all printing layouts (selected in blue). B) Edition of barcode field to activate the newly installed IDAutomationHC39M font. Once the field is selected, just change the font and exit layout saving the changes. C) Photograph of labelled rack. D) A magnified view of the a few tanks and one row of a rack. Note the individual labels on the tanks, which contain relevant information (Tank Unique ID, age, sex, genotype, lab owner, IACUC protocol, and position), as well as the labels on the rack itself, which correspond to the fixed positions/configuration of that rack. Also note the unique barcodes on each tank, and the rack, which facilitate easy identification and moving of tanks, as well as setting up crosses or other functionality.

**Supplemental Figure 3. Creating dead fish reports.** A) “Tank List View” layout with arrow pointing to “Print Dead Fish Report”. B) Example of a weekly dead fish report (that can be printed out).

**Supplemental Table 1. Comparison between FileMaker Server and FileMaker Server Cloud Applications.**

