## Supplementary material for "FishNET: An automated relational database for zebrafish colony management"

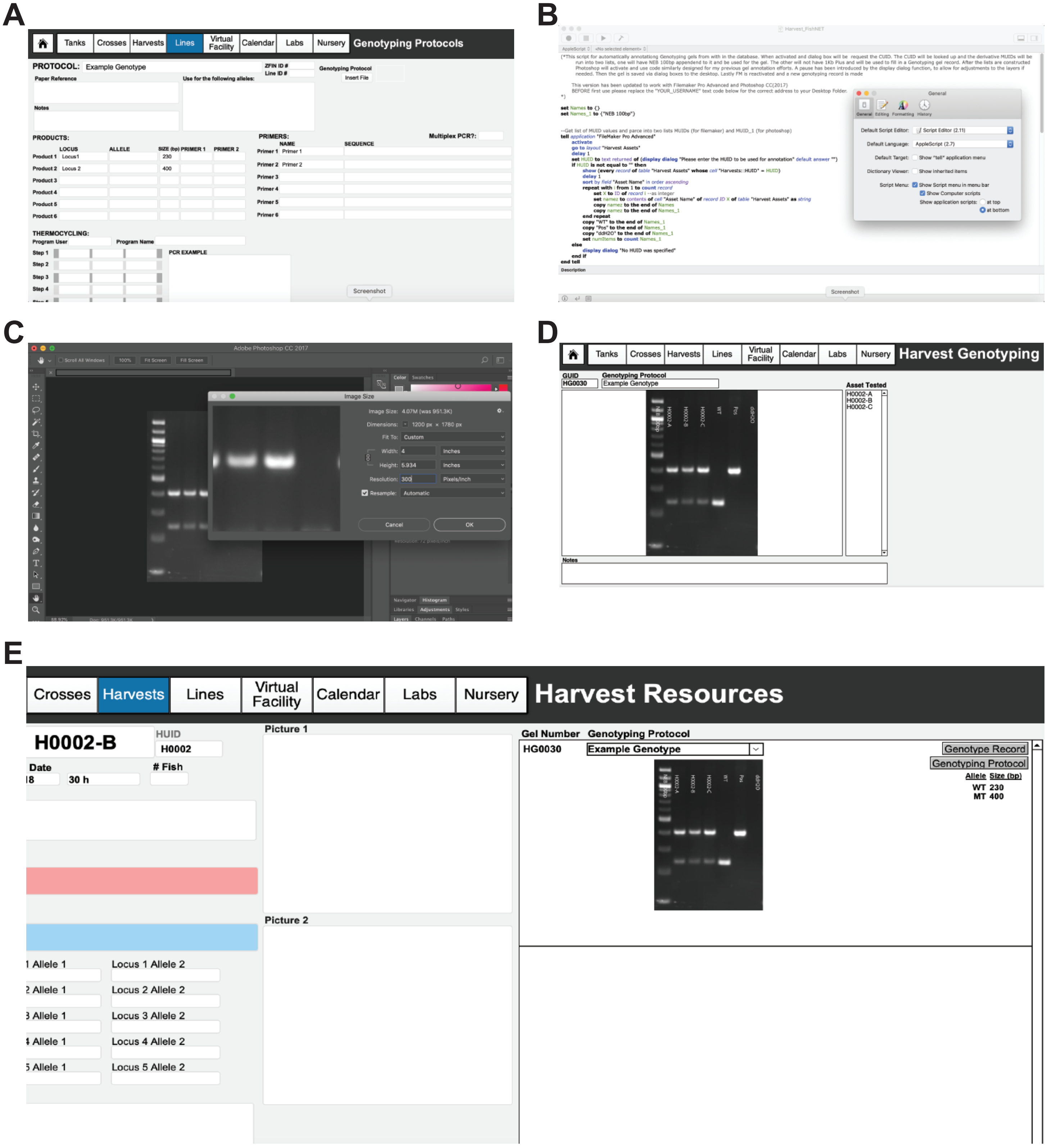


### Supplemental Figure 1

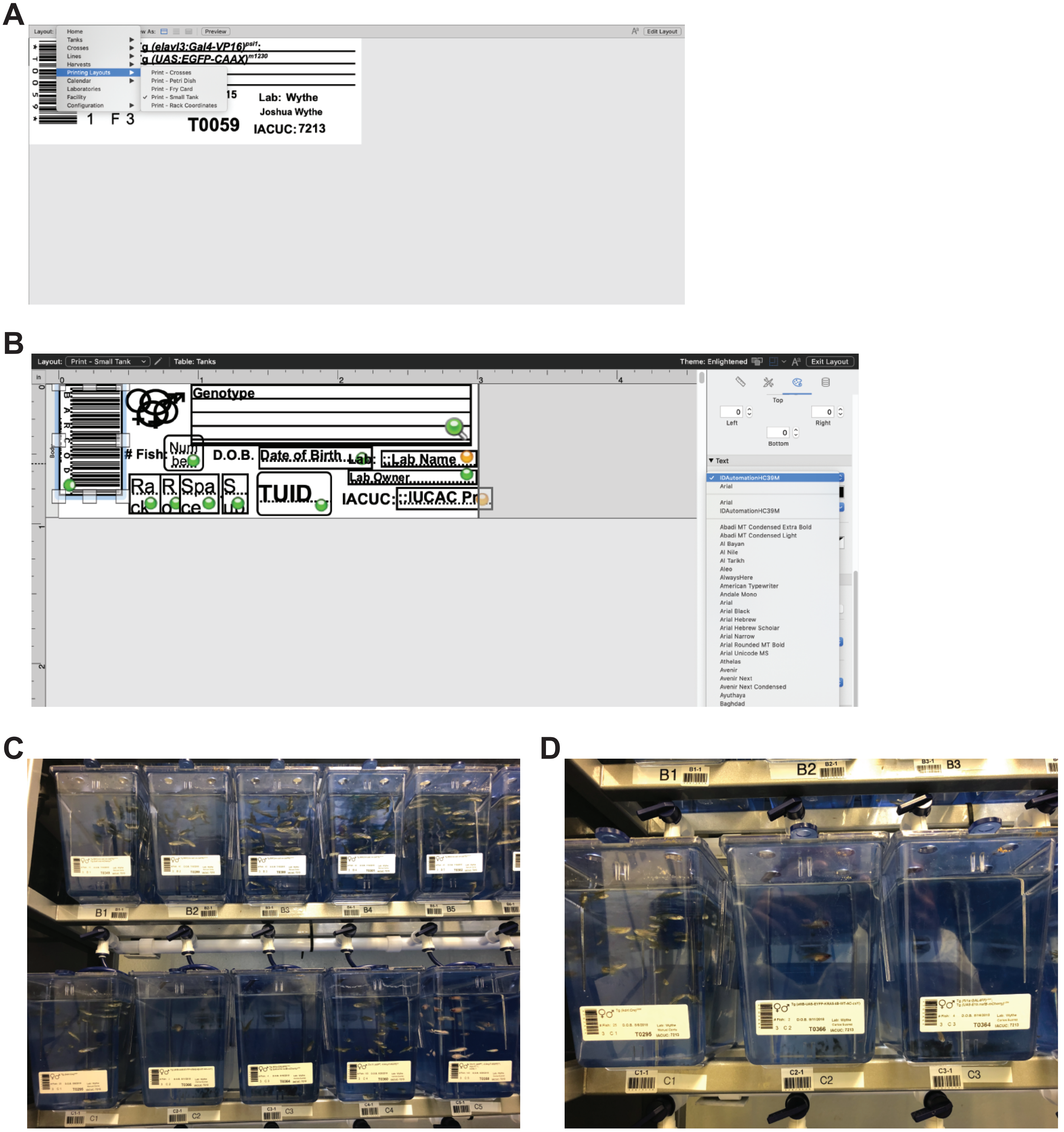


### Supplemental Figure 2

A

B

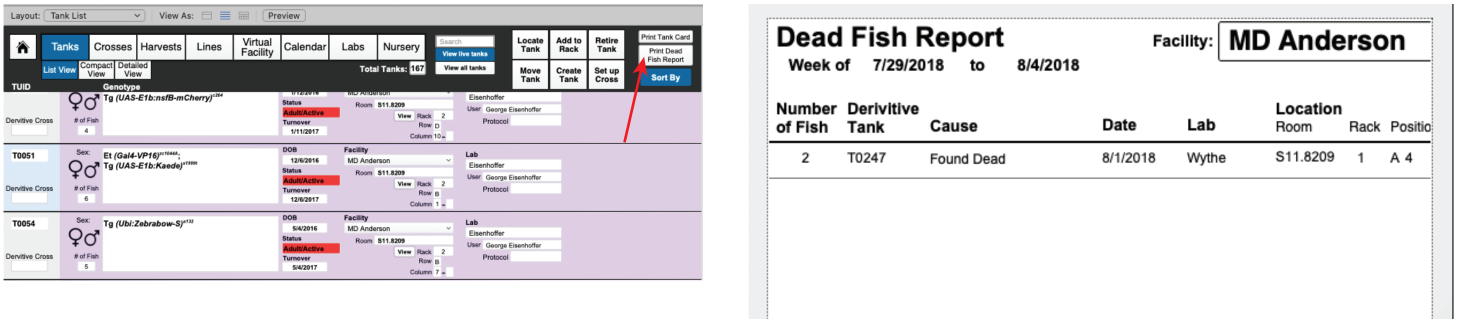


### Supplemental Figure 3

#### Supplemental Table 1. Comparison between FileMaker Server and Cloud Applications

| Category | FileMaker Server | FileMaker Cloud |
|---|---|---|
| **Hosting** | On user premise. | Amazon Web Services (AWS) hosted in the cloud. |
| **Hardware costs** | Server-class hardware required (upgrades and maintenance). | No up-front hardware costs. |
| **Licensing** | Requires either an annual or permanent FMP software license. | Requires an hourly or annual FileMaker software license, plus an AWS subscription that includes services (computing, storage, data transfer, and email) |
| **Capacities** | Tested to support up to 500 FMP Pro Advanced, FMP Go, or FMP WebDirect clients. | Tested to support up to 100 FMP Advanced, FMP Go, or FMP WebDirect clients |
| **Scalability** | May need to buy additional hardware and spend time with setup and configuration. | Quickly scales up or down for seasonal demand periods. |
| **Internet Connection Speed** | Data can be accessed via private data centers and across private LAN connections. | Access to data is dependent on an Internet connection or WAN network. |
| **IT impact** | Requires someone to perform administrative tasks. | Minimal impact to existing technical staff. |
| **Maintenance** | Monitoring and OS updates must be scheduled. | Monitor live status and get automatic notifications for OS updates and software patches. |
| **Backups / Recovery** | Need to create and manage backup schedules. Any backup can be used to recover data. | Backups are created and preserved automatically when auto-maintenance is enabled. |
| **Authentication** | Supports external authentication via Active Directory, Open Directory, and OAuth 2.0 identity providers. | Supports custom app authentication via FileMaker user accounts and OAuth 2.0 identity providers. |
| **Security/ Certificates** | You are responsible for the physical security of your server hardware. | AWS is responsible for the physical security of the server hardware. |
